# Efficient neural encoding of time intervals in speech and complex sound sequences

**DOI:** 10.64898/2026.06.01.729227

**Authors:** Honghua Chen, Jing Wang, Jiaxin Gao, Hang Zhang, Nai Ding

## Abstract

Encoding time intervals in complex sound faces dual challenges: it must be precise and cover a broad dynamic range. In speech, for example, a ten millisecond lengthening of a syllable can signal stress or phrasal boundaries, yet the syllable duration distribution is long tailed beyond 500 ms and has variable statistics across speakers. Here, we propose that the auditory cortex employs efficient coding to represent time intervals. When listeners heard syllable sequences drawn from different duration distributions, the magnetoencephalographic (MEG) response from the temporal cortex scaled with syllable duration, which is characterized using the interval-response function. Crucially, this interval–response function met predictions of efficient coding. Its intercept and slope adapted to the mean and variance of syllable duration, respectively, and it consistently exhibited a compressive nonlinearity that reduced response skewness, consistent with a maximum-entropy code. A computational model that constantly updates the inference of duration distribution provided an algorithmic account of this efficient coding, and intracranial electroencephalogram (iEEG) data confirmed the same principles during natural speech comprehension. Together, our findings reveal an efficient neural mechanism that supports precise encoding of highly variable time intervals in complex sound sequences.

## INTRODUCTION

Timing provides fundamental information for making sense of complex sound sequences, including speech, music, and other natural sounds. For example, in speech perception, a 10–20 millisecond lengthening of syllable duration can signal a word or phrase boundary (Blum et al., 2024; Paschen et al., 2022) or convey communicative intent such as emphasis (Katz & Selkirk, 2011; Kügler, 2008). Similarly, in music, notes played a few tens of milliseconds off the beat can be perceived as syncopation and induce musical groove (Harding et al., 2025; Vuust et al., 2022). Critically, the demand for precise time encoding co-exists with the demand to cover a broad dynamic range of time intervals. For example, in “a scratch”, the syllable “a” may last less than 50 ms while the syllable “scratch” may last nearly 500 ms. In music, a sixteenth note is 1/16 of the duration of a whole note. Since the brain operates under tight metabolic constraints, precisely encoding such a wide range of intervals requires an efficient code (Barlow, 1961; Barlow & Levick, 1969; Brenner et al., 2000; Laughlin, 1981). Previous studies have identified candidate neural codes for time intervals (Buzsáki, 2026; Kösem et al., 2018; Meirhaeghe et al., 2021; Paton & Buonomano, 2018; Tsao et al., 2022). For example, the field potential evoked at the end of a time interval scales with the duration of the interval (Kononowicz & Rijn, 2014; Ofir & Landau, 2022; Wiener & Thompson, 2015; Y. Zhang et al., 2026). However, it remains unclear whether the neural code of time intervals is optimized for efficiency.

According to the efficient coding theory, the most efficient neural code is one that adapts to the input statistical distribution (Młynarski & Hermundstad, 2021; Weber et al., 2019). Such adaptation can occur on two time scales. On a shorter timescale, stimulus-specific statistics may drive adaptation, and previous animal studies have confirmed rapid adaptation to the mean and variance of sound or light intensity (Carandini & Heeger, 2012; Dean et al., 2005; Rabinowitz et al., 2011; Schwartz & Simoncelli, 2001). On a longer timescale, adaptation to statistical regularities that are stable across natural stimuli may be hardwired through evolution. For example, previous studies have shown that the auditory and visual receptive fields can be explained as optimal encoders for natural sounds or images, suggesting evolutionary adaptation (Gervain & Geffen, 2019; Lewicki, 2002; Pitkow & Meister, 2012; Smith & Lewicki, 2006; Soto et al., 2020).

When applied to the neural encoding of time intervals in sound, efficient coding theory predicts adaptation to interval distributions, and such adaptation may also occur on two time scales. On the one hand, in natural sound, the mean and variance of time intervals are highly variable and stimulus-specific. For example, mean syllable duration differs widely across speakers (Jacewicz et al., 2009) and languages (Coupé et al., 2019; Pellegrino et al., 2011; Y. Zhang et al., 2023), and musical tempo also differs widely across songs (Mehr et al., 2019; Passmore et al., 2024). Furthermore, previous behavioral and neural studies have demonstrated adaptation to mean time intervals (Dilley & Pitt, 2010; Kösem et al., 2018; Maslowski et al., 2019; Reinisch & Sjerps, 2013). For instance, a vowel is more likely to be perceived as a long vowel when the mean syllable duration is shorter. On the other hand, the interval distributions tend to be positively skewed, which is characterized by a majority of short intervals and a small number of long intervals, creating a long right tail. For example, the distribution of speech syllable duration is demonstrated to be positively skewed (Greenberg et al., 2003; Y. Zhang et al., 2023). Although this phenomenon has to be confirmed for other sound categories, theoretically, the time interval between two random events, such as two raindrops, is subject to the exponential distribution, which is strongly positively skewed. The two-timescale framework of efficient coding therefore predicts that the neural code for time intervals may rapidly adapt to the mean and variance of the interval distribution, while exhibiting hardwired optimization for positively skewed distributions.

Here, we tested these predictions by investigating whether and how the neural code for time intervals adapts to the mean, variance, and skewness of interval distributions. We first analyzed the statistical distribution of time intervals in natural sound and confirmed that intervals are generally positively skewed, while the mean and variance are highly variable. Next, we conducted three MEG experiments on synthesized syllable sequences. The three experiments separately manipulated the mean, variance, and skewness of the interval distribution and analyzed the neural response evoked at the offset of each interval (Kononowicz & Rijn, 2014; Ofir & Landau, 2022; Wiener & Thompson, 2015; Y. Zhang et al., 2026). We then built a computational model to account for the observed adaptation effects. Finally, guided by the model, we analyzed two intracranial EEG (iEEG) datasets to test whether the neural code identified with controlled stimuli generalizes to natural speech comprehension.

## RESULTS

### Statistics of Time Intervals in Speech and Natural Sound

We characterized the distribution of syllable duration in natural speech based on six speech corpora that spanned two languages (Chinese and English) and three speaking styles (reading sentences, reading audiobooks, and spontaneous speech; Fig. 1A-B). The syllable duration widely varied across corpora and speakers (Fig. 1C-E). The mean syllable duration of an individual speaker varied between 156 and 262 ms (5th–95th percentile), and the standard deviation varied between 58 and 127 ms (5th–95th percentile). The distributions were consistently positively skewed across speakers (5th–95th percentile of skewness was between 0.38 and 1.77). We also characterized the interval distribution of other representative natural sound sequences, including human song, birdcall, and human footstep (Fig. 1F). The note durations in human song, syllable durations in birdcall, and step intervals in footstep exhibited distinct timescales, but their skewness was consistently positive (Fig. 1G). These results revealed that the distribution of time intervals in natural sound was generally positively skewed, while the mean and variance were highly variable.

**Figure 1:**
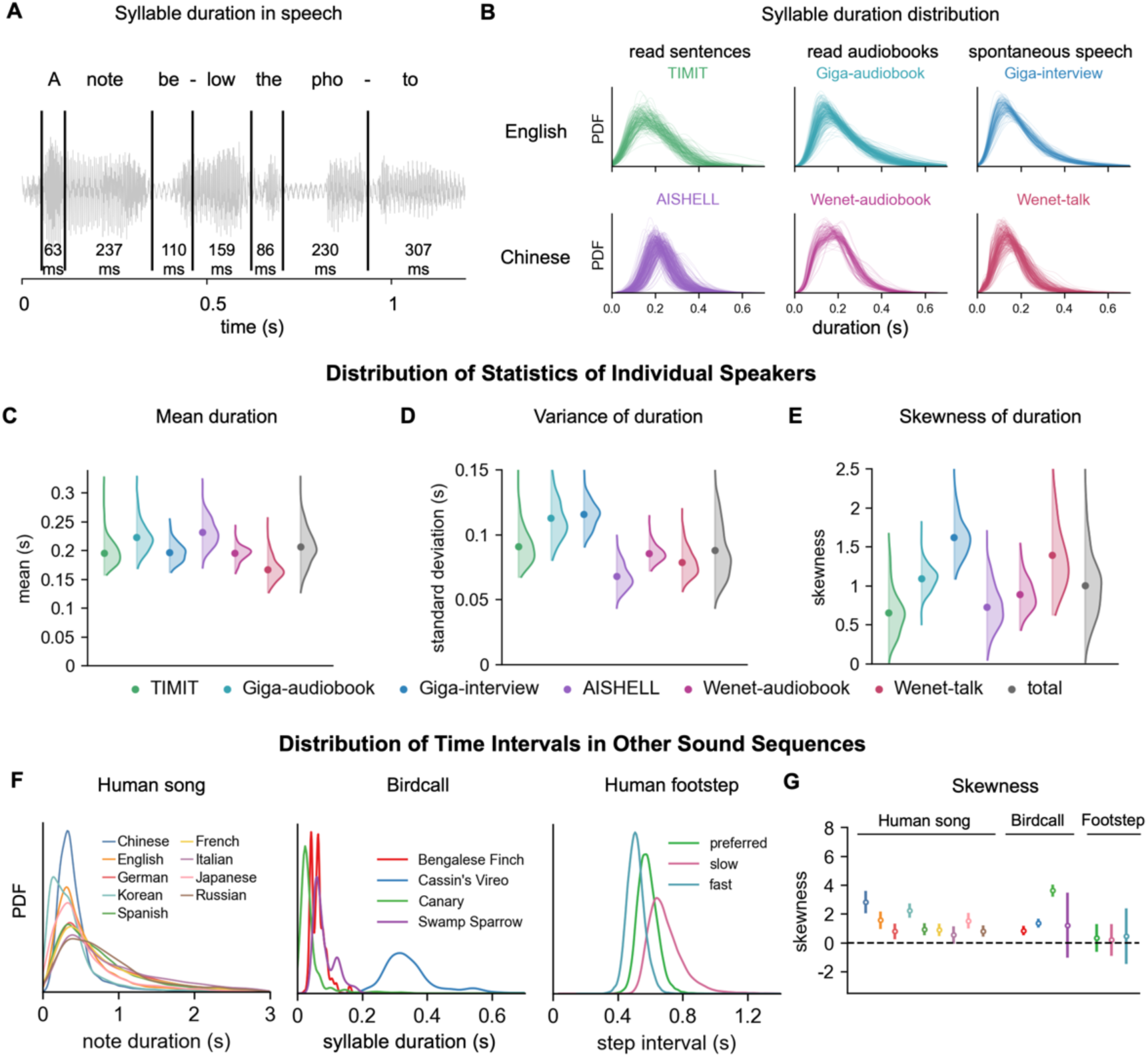
Statistics of time intervals in natural speech. (A) Example English speech with marked syllable time intervals. (B) Distribution of syllable duration for individual speakers across six corpora spanning different languages and speaking styles. (C-E) For recordings of each speaker, we calculated the mean, variance, and skewness of syllable duration. The distribution of the mean, variance, and skewness was shown. Each dot indicates the mean statistic across speakers. (F) Distributions of time intervals for human songs in nine languages, birdcall of four species, and human footsteps. (G) Skewness of time intervals for distributions shown in panel F (same color for each condition). Skewness was calculated for each person or bird. Error bars indicate the SD of the skewness distribution.

### Efficient coding hypothesis

According to the efficient coding theory (Barlow, 1961; Laughlin, 1981), the neural system maximizes the response entropy to efficiently encode stimulus information (Fig. 2A). When the neural response power is constrained, the response that most efficiently encodes the stimulus is subject to a Gaussian distribution, i.e., the maximum-entropy distribution (Cover & Thomas, 1999), regardless of the stimulus distribution. Specifically, the Gaussian distribution has zero mean and its variance equals the power constraint. To transform an arbitrary stimulus distribution into the desired Gaussian distribution, the neural system must nonlinearly scale the stimulus according to the stimulus distribution. Here, we refer to this nonlinear scaling function that links the stimulus interval and neural response as the interval-response function. The efficient coding theory generates three testable predictions. First, as the stimulus mean increases, the x-axis intercept of the interval-response function should shift rightward to maintain a constant response mean (Fig. 2B). Second, as the stimulus variance decreases, the gain (slope) of the interval-response function should increase to preserve the response variance (Fig. 2C). Third, when the stimulus distribution is skewed, the interval-response function should exhibit a nonlinearity that reduces the response skewness toward zero (Fig. 2D). Simulation analysis demonstrates the efficiency of these adaptation strategies (Fig. S1). In the following, we test these three predictions using three MEG experiments, build a computational model to implement efficient interval coding, and verify the model using two iEEG datasets.

**Figure 2:**
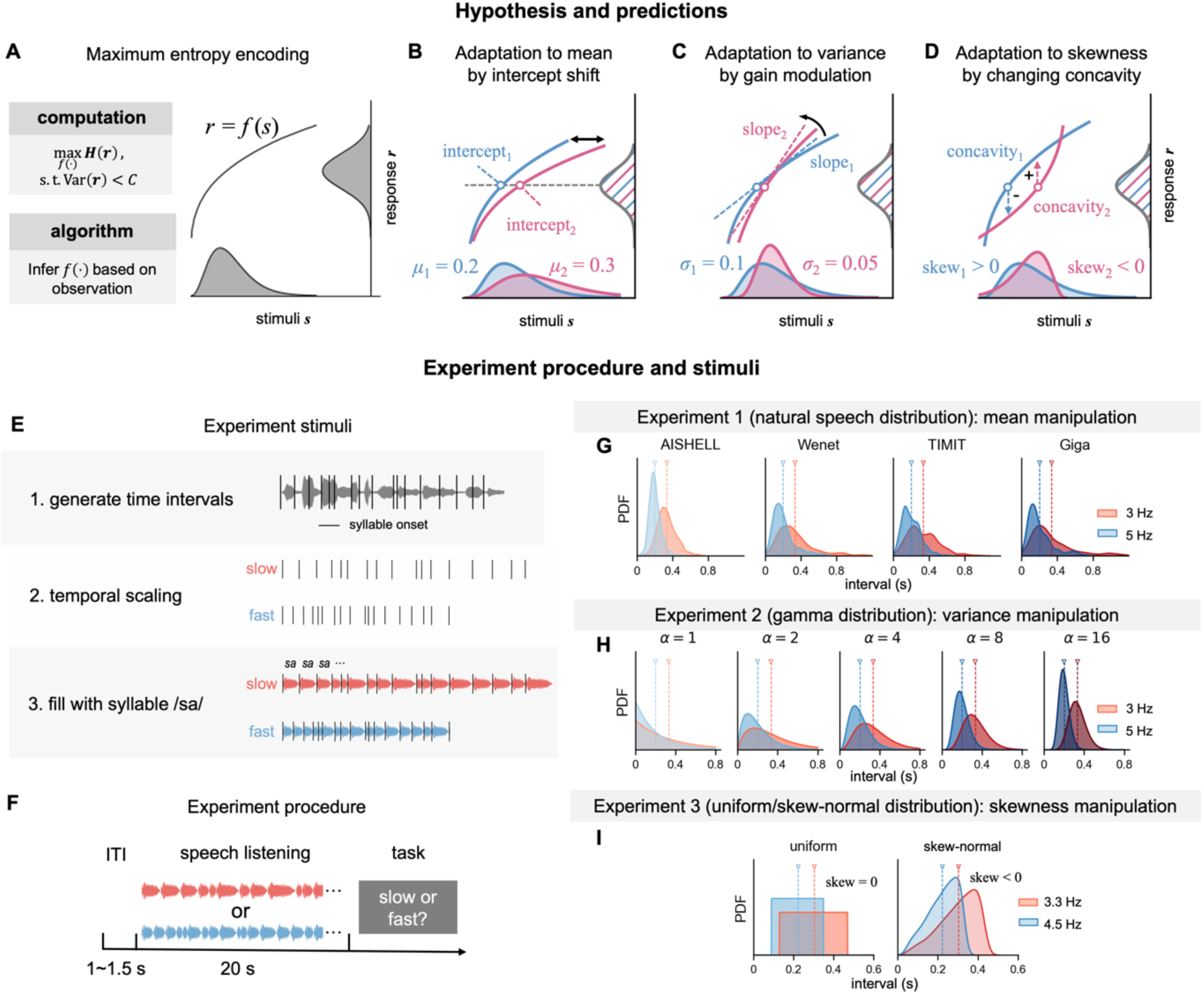
Hypothesis and experiment design. (A) Schematic of the efficient coding of time intervals. The neural system infers a nonlinear function, i.e., the interval-response function, based on the observed stimulus distribution to maximize the entropy of neural response. (B–D) Predictions of efficient neural coding of time intervals. (B) The interval-response function adapts to the mean duration by shifting the x-axis intercept. (C) The interval-response function adapts to the duration variance by modulating the gain (slope). (D) The interval-response function adapts to skewness by adjusting the concavity to drive the response skewness toward zero. (E) Construction of stimulus. Syllable onsets were extracted from natural speech (Experiment 1) or sampled from parametric distributions (Experiments 2 and 3), temporally scaled to two rates, and filled with the syllable /sa/ (see Methods). (F) Experimental procedure. In each trial, participants listened to a 20-s syllable sequence and the task was to distinguish fast and slow sequences. (G-H) Time interval distributions of the stimulus in Experiments 1–3. (G) Syllable duration extracted from natural speech recordings from four corpora and temporally scaled to the mean syllable rate of 3 Hz or 5 Hz. (H) Each syllable duration independently drawn from Gamma distributions with different shape parameters *α*, which manipulate variance independently of the mean. (I) Each syllable duration independently drawn from uniform and skew-normal distributions that manipulate skewness while maintaining mean and variance.

### MEG experiments

In three MEG experiments, we constructed sequences based on a single syllable /sa/ and manipulated the duration of each syllable. Compared with natural speech, in the synthesized sequences, all syllables had the same phonetic content and only varied in duration. Experiments 1, 2, and 3 separately manipulated the mean, variance, and skewness of the syllable-duration distribution. In Experiment 1, the syllable duration sequence was extracted from real speech recordings and was temporally scaled to manipulate the mean syllable rate, i.e., the average number of syllables per second (Fig. 2G). In Experiment 2, the syllable duration was drawn from Gamma distributions, and the standard deviation of duration was explicitly manipulated using the shape parameter of the Gamma distribution (Fig. 2H). In Experiments 1 and 2, the duration distributions were positively skewed, consistent with the statistical properties of natural sound. Experiment 3, however, tested syllable-duration distributions that had zero or negative skewness using uniform and skew-normal distributions, respectively.

In all experiments, the stimuli were separated into a fast condition and a slow condition, and the participants’ task was to judge whether the sequences were fast or slow (Fig. 2E). Behavioral accuracy was high in all experiments (96.2% ± 3.1%, 93.4% ± 5.4%, and 90.0% ± 5.9% for Experiments 1, 2, and 3, respectively). We recorded magnetoencephalography (MEG) responses while participants listened to the sequences and analyzed the neural responses from the 204 gradiometers. To isolate neural responses that encode time intervals, we regressed out neural responses to basic acoustic features using the temporal response function (TRF) before further analysis (see Supplementary Information).

### Neural Encoding of Syllable Duration

We first investigated whether a neural response encoding syllable duration could be observed at the offset of each syllable, as was suggested by earlier studies (Kononowicz & Rijn, 2014; Ofir & Landau, 2022; Wiener & Thompson, 2015; Y. Zhang et al., 2026). The syllable offset was the earliest moment when the syllable duration could be determined. We extracted the MEG response to each syllable and time-aligned them based on the syllable offset. We quantified the cross-validated correlation between neural response and syllable duration at each time lag relative to the syllable offset, for each MEG channel (Fig. 3A; see Methods). This time-resolved correlation averaged across all conditions showed a peak at approximately 100 ms after syllable offset, in channels over temporal regions (Fig. 3A, inset; permutation t-test with FDR correction, P < 0.05). In other words, about 100 ms after a syllable ended, we observed a neural response reflecting how long the syllable lasted. To further validate the effect, we visualized the neural response to syllables of different durations and confirmed that these responses clearly diverged about 100 ms post syllable offset (Fig. 3B). The scatter plot between syllable duration and neural response at peak latency (Fig. 3C) suggested a nonlinear relationship between the two. This nonlinear function was better fitted by a sigmoidal function than a linear or logarithmic function (Fig. 3D; model fitting using cross-validated correlation, with model comparison using two-sided t-test with FDR correction, P < 0.001). The sigmoidal fit was referred to as the interval-response function in the following analyses. These encoding results were consistent for each condition (Fig. 3EF) and generalized to Experiments 2 and 3 (Fig. S2), establishing a robust neural marker encoding time intervals.

**Figure 3:**
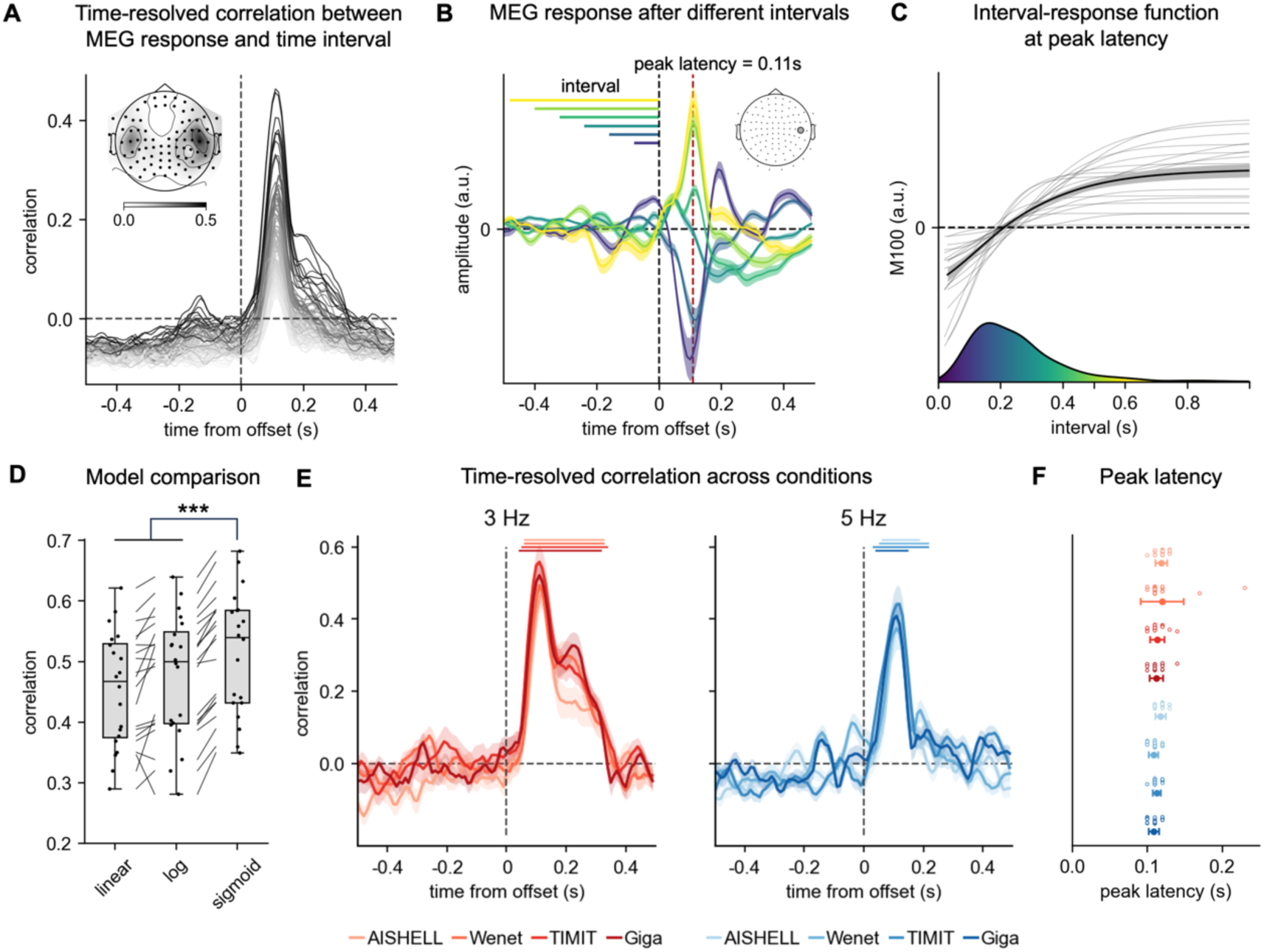
Neural encoding of time intervals. (A) Time-resolved correlation between MEG response and syllable duration. Each curve is the response from a MEG channel, and only responses from channels showing a significant correlation with syllable duration are shown (i.e., channels shown by dots in the topography, P < 0.05, permutation t-test with 5,000 permutations, FDR-corrected). (B) Time course of neural activity for syllables binned by time interval, from a representative channel. Shading indicates SEM of participants. (C) Interval-response function at peak latency. Gray lines show individual participants and the black line shows the average across participants. The histogram below shows the stimulus interval distribution. Shading indicates SEM across participants. (D) Interval-response function fitted by three models. Whisker plots show the distribution of correlation at peak latency for each model (center line, median; box, 25th–75th percentile; whiskers, ±1.5 × interquartile range; dots, individual participants). ***P < 0.001, two-sided t-test with FDR correction. (E) Time-resolved correlation between MEG response and syllable duration for each 3-Hz (left) or 5-Hz (right) condition. Shading indicates SEM of participants. Horizontal bars on top denote significant correlation (cluster-based permutation test, P < 0.05; cluster-forming using point-wise t-test, P < 0.01 with 5,000 permutations). (F) Peak latency of encoding correlation for all conditions. Error bars indicate SD across participants.

### Adaptation to Duration Mean

We next tested the three hypotheses illustrated in Fig. 2 by analyzing the interval-response function. Specifically, we focused on the interval-response function estimated based on the neural response at the latency that maximized the correlation between neural response and syllable duration, which was consistently distributed between 100 ms and 120 ms. We averaged five peak channels as regions of interest (ROI) to get the interval-response function for each participant. In Experiment 1, consistent with prediction 1, the interval-response function in the 5-Hz conditions consistently shifted leftward compared with the interval-response function in the 3-Hz conditions (Fig. 4A): The intercept of the interval-response function (averaged over four corpora) was significantly smaller in 5-Hz conditions (two-sided t-test, P < 0.001). Separation of interval-response functions between fast and slow rates was also observed in Experiments 2 and 3 (Fig. 4DG): The intercept of the interval-response function was significantly smaller in the fast conditions (Fig. 4EH; two-sided t-test, P < 0.001).

**Figure 4:**
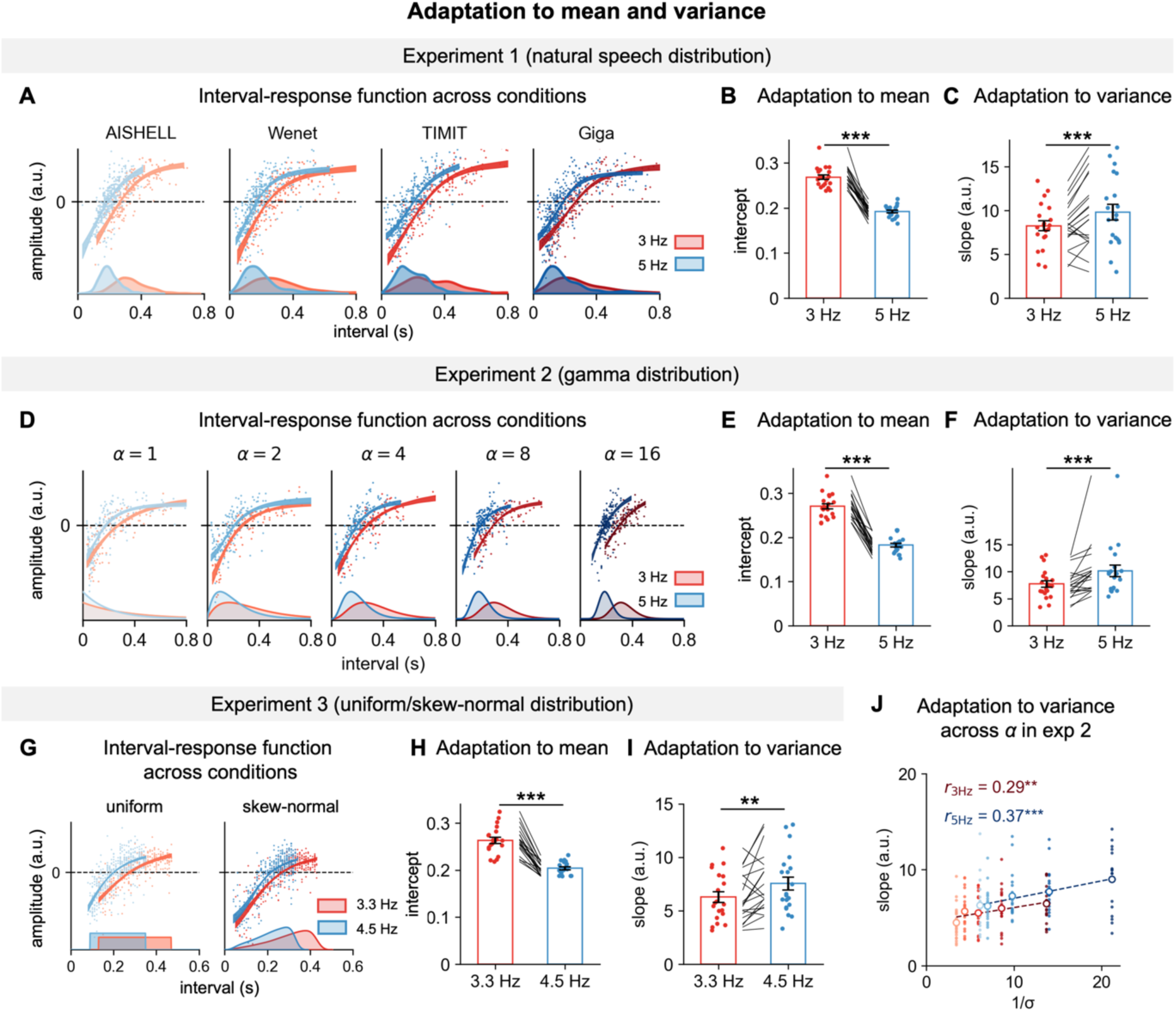
Adaptation to mean and variance of time interval. (A) Interval-response functions in Experiment 1. Shadow around the curve shows the s.e.m. across participants. Each dot was the neural response to each syllable (*N* = 240 for 3-Hz conditions, *N =* 400 for 5-Hz conditions), averaged over participants. Stimulus interval distributions are shown below. (B, C) Intercept and slope of interval-response functions. **P < 0.01, ***P < 0.001, two-tailed paired-sample t-test. (D–F) Same as (A–C) for Experiment 2. (G–I) Same as (A–C) for Experiment 3. (J) Relationship between the slope of interval-response function and the reciprocal of the standard deviation of stimulus intervals. All conditions in Experiment 2 are shown (same color as panel D), which differ in the shape parameter of Gamma distribution. **P < 0.01, ***P < 0.001, Pearson correlation test.

### Adaptation to Duration Variance

In Experiment 1, the duration variance also differed between 3 Hz and 5 Hz, as scaling of the mean syllable rate also scaled the variance. Consistent with prediction 2, the interval-response function exhibited a steeper slope in 5-Hz conditions that had smaller variance (Fig. 4C; two-sided t-test, P < 0.001). In other words, the interval-response function in the 5-Hz condition had a higher gain that compensated for the decrease in duration variance. The slope change of the interval-response function between fast and slow conditions was also observed in Experiments 2 and 3 (Fig. 4FI; two-sided t-test, P < 0.001 in Experiment 2 and P < 0.01 in Experiment 3). Furthermore, to dissociate the effect of variance change and the effect of mean change, we separately manipulated variance at each mean syllable rate in Experiment 2. Specifically, the time intervals were sampled from gamma distributions with different shape parameters (α = 1, 2, 4, 8, 16), which could manipulate the variance of the distribution independent of the mean (Fig. 4D). Adaptation to variance was also observed when the stimulus variance varied within each mean syllable rate (Fig. 4J). The slope of the interval-response function scaled with the reciprocal of the standard deviation (Pearson’s r = 0.29, P < 0.01 for 3 Hz; Pearson’s r = 0.37, P < 0.001 for 5 Hz), consistent with the prediction of adaptation to variance.

Across all three experiments, adaptation to mean and variance was predominantly localized to the right temporal cortex (Fig. S3). Together, these results demonstrate that the brain adapts to both the mean and variance of the temporal distribution, matching the dynamic range of neural responses to stimulus statistics in a manner that enhances encoding efficiency.

### Adaptation to Duration Skewness

Next, we investigated whether the interval-response function exhibited nonlinearity that transformed skewed input distributions toward a zero-skewness Gaussian response distribution (Fig. 2D). We first quantified whether the skewness of the MEG response was closer to zero than the skewness of the stimulus. Since the MEG measurement was mixed with noise that could be assumed to be Gaussian, we controlled the noise level of the response when quantifying its skewness (see Methods).

In Experiment 1, the skewness of MEG response was significantly reduced compared with the skewness of stimulus syllable duration (Fig. 5A; two-tailed paired-sample t-test, P < 0.001 for all conditions). The absolute value of the skewness was also significantly reduced in the MEG response than in the stimulus in three of the four conditions (two-tailed paired-sample t-test, FDR corrected, P < 0.001 for syllable duration drawn from the Wenet, TIMIT, and Giga datasets). In Experiment 2, the skewness of the MEG response was significantly reduced compared with the stimulus (Fig. 5B; two-tailed paired-sample t-test, P < 0.001). The absolute value of the skewness was reduced when the stimulus gamma distribution was highly skewed, i.e., *α* = 1 or 2 (two-tailed paired-sample t-test, FDR corrected, P < 0.001). However, when the stimulus intervals were drawn from a relatively symmetric gamma distribution (i.e., *α* = 8 or 16), the absolute skewness of the MEG response was increased compared with the stimulus (two-tailed paired-sample t-test, P < 0.001). Similarly, in Experiment 3, when the stimulus distribution was symmetric or negatively skewed, the MEG response was more negatively skewed than the stimulus (Fig. 5C; two-tailed paired-sample t-test, P < 0.001). Lastly, the second derivative of the interval-response function was significantly negative (Fig. 5D; two-tailed paired-sample t-test, P < 0.001), indicating that the interval-response functions across all conditions exhibited compressive nonlinearity (Fig. 3). These results suggested that the brain employed a stereotypical compressive nonlinearity that reduced response skewness. This nonlinearity was well-suited for efficiently encoding positively skewed distributions, which are commonly found in natural sound (Fig. 1).

**Figure 5:**
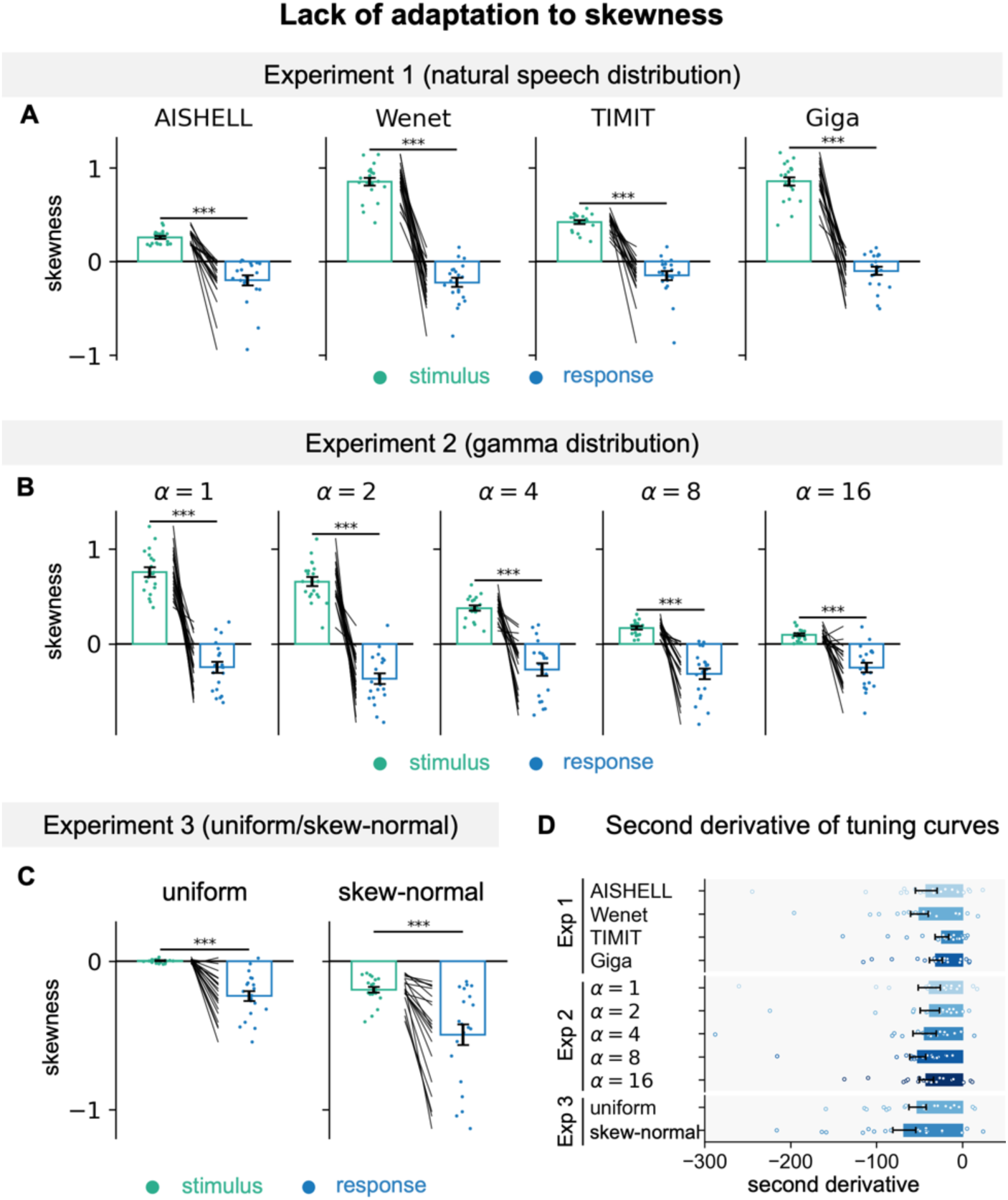
Response nonlinearity efficiently encodes positively skewed interval distributions. (A) Skewness of stimulus time intervals and skewness of the neural response in Experiment 1. For the neural response, noise levels were calibrated based on inter-trial correlation (ITC) of neural activity (see Methods). ***P < 0.001, two-tailed paired-sample t-test. (B, C) Same as (A) for Experiments 2 and 3. (D) Second derivative of the interval-response function across all conditions in Experiments 1-3. Negative second derivative indicates that the interval-response function shows compressive nonlinearity.

### Modeling Efficient Neural Coding of Time Intervals

The previous analyses established that the neural response flexibly adapted to the mean and variance of stimulus interval distribution. Next, we modeled how the brain might infer the interval-response function online based on the intervals in each trial (Fig. 6A). The model calculated running estimates of mean and variance of intervals using an exponentially weighted moving average (EWMA). These estimates were mapped to the intercept and slope of the interval-response function using linear transforms. Parameters of these linear transforms, as well as the learning rate of EWMA, were fitted for each participant based on all stimulus conditions.

**Figure 6:**
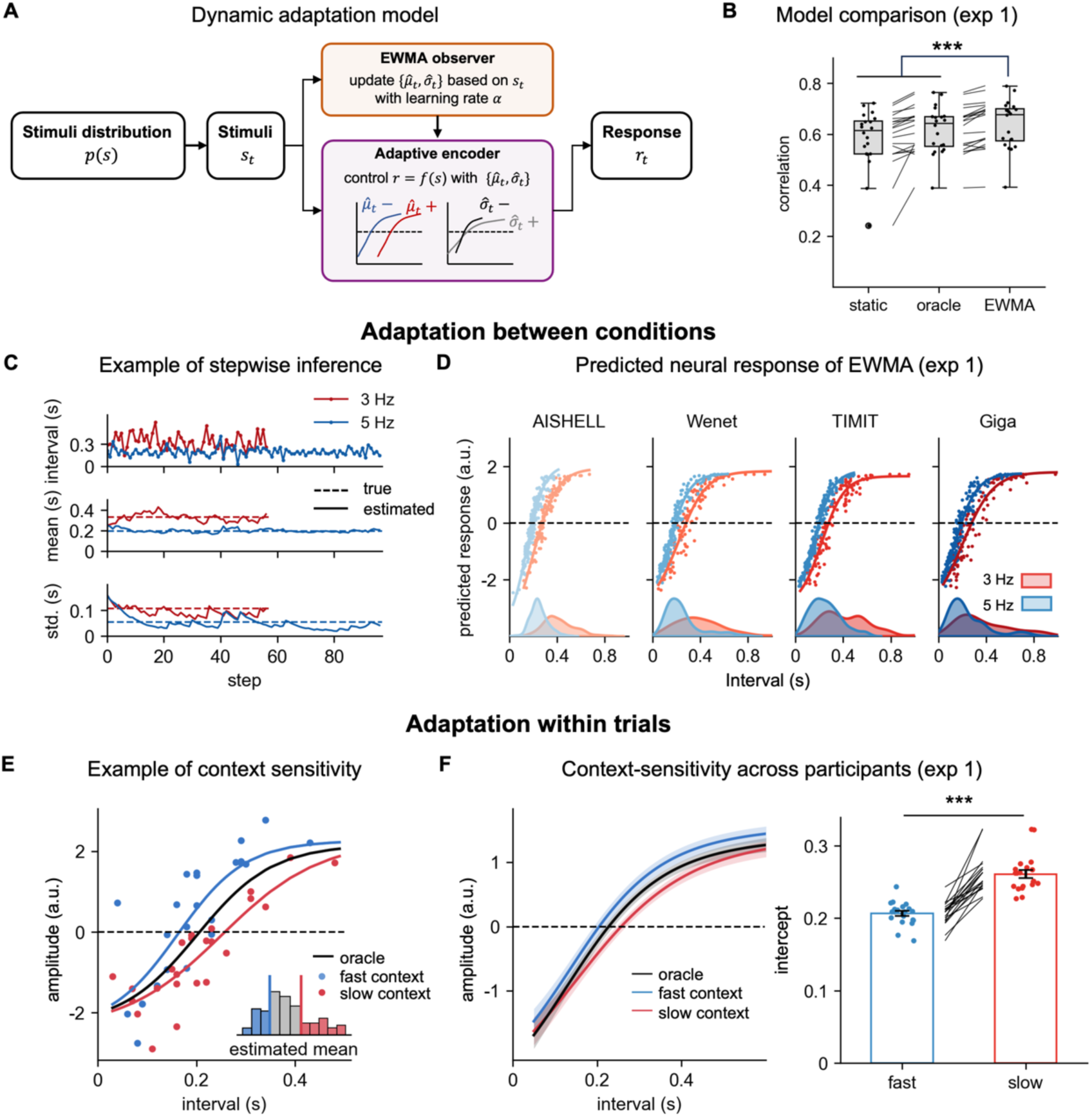
Modeling neural adaptation to interval mean and variance. (A) Schematic of the model. The observer iteratively infers the mean and variance of the stimulus distribution using an exponentially weighted moving average (EWMA) with a fixed learning rate. The adaptive encoder uses these estimated statistics to update the intercept and slope of the interval-response function. (B) Model comparison. Three models with different context estimation strategies were fitted to the data (see Methods). Box plots show cross-validated correlations (center line, median; box, 25th–75th percentile; whiskers, ±1.5 × interquartile range). The EWMA model outperforms all alternatives (***P < 0.001, two-sided t-test with FDR correction). (C) Example inferred mean and standard deviation (one participant in Experiment 1). (D) Model-predicted interval-response functions across conditions for the same participant in panel C. Scatter plots show model predictions of neural activity; lines show the fitted static interval-response function for model predictions. (E) Context-dependent MEG responses. Dots show intervals and corresponding M100 responses, classified by estimated EWMA mean (inset: threshold at 25th/75th percentile, defining fast and slow contexts). Corresponding interval-response functions were fitted separately for fast and slow contexts; the oracle interval-response function was fitted to all intervals. (F) Context sensitivity across participants in Experiment 1. Left: Interval-response functions averaged across conditions and participants; shading indicates s.e.m. of participants. Right: Intercepts of the interval-response functions for fast and slow contexts. ***P < 0.001, two-tailed paired-sample t-test.

The EWMA model better fitted the MEG responses than a static model that used a fixed interval-response function, as well as the oracle model that had perfect knowledge of each trial’s statistics (Fig. 6B; model fitting using cross-validated correlation, with model comparison using two-sided t-test with FDR correction, P < 0.001). The EWMA also outperformed other alternative models, e.g., a model with only mean or variance estimation (see Fig. S4 for extended model comparison). Figure 6C illustrates the adaptation process from a participant, in which the EWMA model’s internal estimates of mean and variance evolved continuously as new intervals were encountered. The interval-response function based on the estimated mean and variance reproduced the adaptation effects to mean and variance observed in the MEG experiments (Fig. 6D). The learning rate of the EWMA indicated the degree of adaptation to recent observations. The fitted learning rate was 0.40 on average (0.46, 0.38, and 0.37 in Experiments 1, 2, and 3, respectively), suggesting rapid adaptation. More importantly, the model revealed adaptation to the within-trial fluctuations of running mean and variance (see Fig. 6E for an example participant). Specifically, when we distinguished locally fast and slow contexts within the same trial based on the estimated running mean, we found the interval-response function shifted leftward in the fast local contexts than in the slow local contexts (Fig. 6F; two-sided t-test, P < 0.001). In other words, the same interval would generate different neural responses depending on whether the preceding intervals were long or short. The same effects were replicated in Experiments 2 and 3 (Fig. S5). These results demonstrated that neural activity adapted to the local context. Since this local adaptation effect could be validated without explicitly manipulating the stimulus mean interval, it could be tested when participants listened to natural speech from a single speaker. Serial dependency analysis further demonstrated the dependence of the MEG response on local histories and validated the EWMA model (Fig. S6).

### Adaptive Time Encoding in Natural Speech Understanding

In the MEG experiments, the mean syllable rate was manipulated to create fast and slow trials, and the task explicitly engaged estimation of the mean syllable rate. We next asked whether efficient neural encoding of time intervals generalized to natural speech comprehension by analyzing low-frequency (i.e., <40 Hz) intracranial electroencephalogram (iEEG) activity recorded from participants listening to natural spoken narratives either in English (Fig. 7) or Chinese (Fig. S7). We found that the field potential in 87 electrodes (out of 1,268 available electrodes) significantly scaled with the duration of the preceding syllable (Fig. 7C), and these electrodes predominantly localized near auditory cortex (Fig. 7E). The interval-response function estimated from each electrode consistently exhibited compressive nonlinearity (Fig. 7D). Critically, the EWMA better explained the iEEG response to syllable duration than the static model (Fig. 7F; two-sided t-test, P < 0.001). Consistently, the interval-response functions estimated in faster or slower local contexts significantly differed in their intercept (Fig. 7G; two-sided t-test, P < 0.001). The serial dependency analysis also validated the dependence of the neural response on local histories and showed the diversity of this dependence across electrodes (Fig. S8). These findings confirmed that the adaptive efficient coding strategy observed in the MEG experiments when listening to artificial stimuli could generalize to naturalistic speech comprehension.

**Figure 7:**
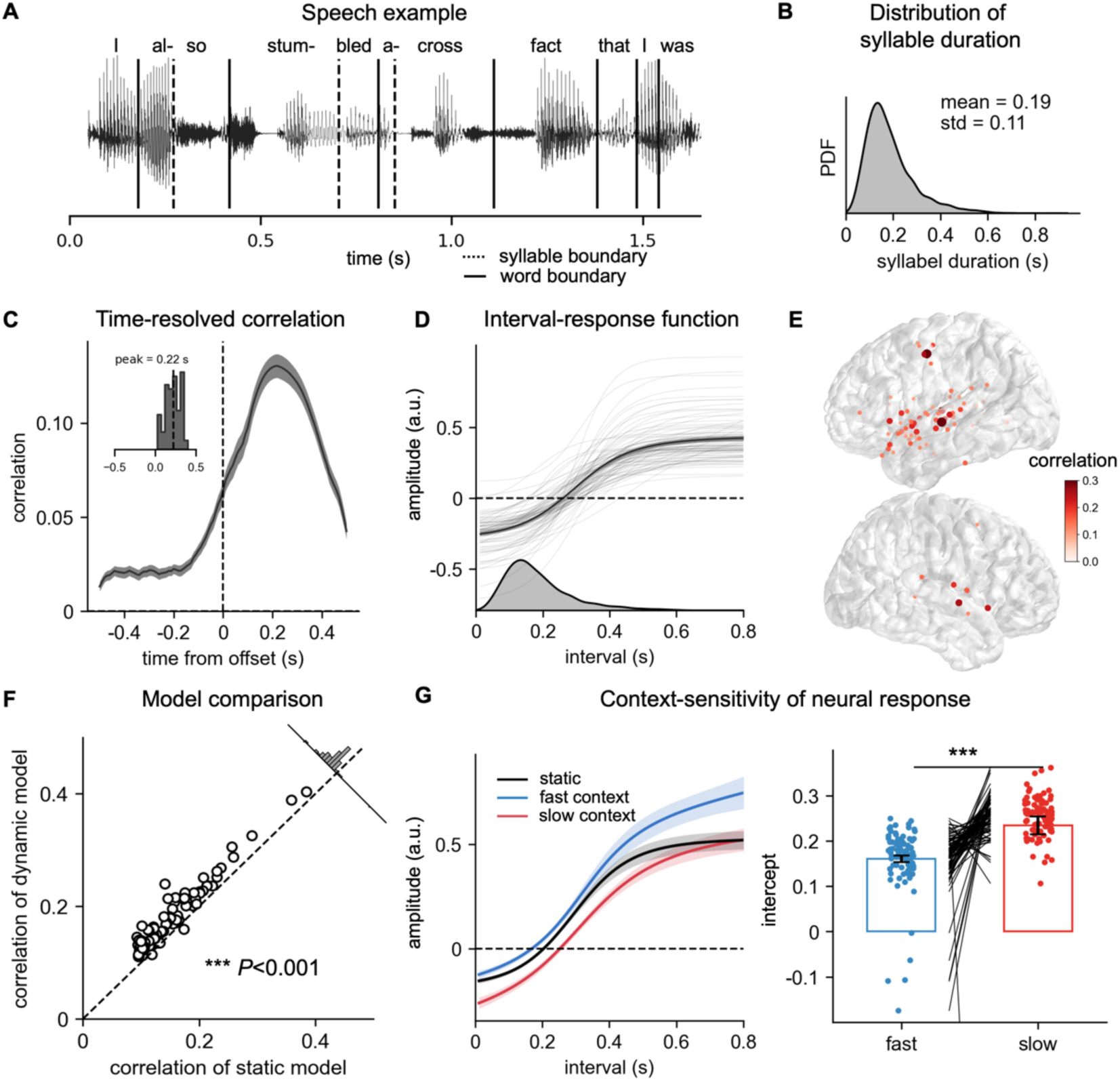
Efficient neural coding of time intervals during natural speech comprehension. (A) Example segment of the speech stimulus. (B) Distribution of syllable duration. (C) Time-resolved correlation of significant electrodes (N = 87). Inset: distribution of peak latency for correlation across electrodes. (D) Interval-response functions for significant electrodes. The histogram below shows the interval distribution. (E) Spatial distribution of encoding strength across electrodes. (F) Model comparison between the dynamic adaptation model and the static sigmoidal model. ***P < 0.001, two-tailed paired-sample t-test. (G) Context-dependent neural responses across electrodes. Left: Interval-response functions for intervals classified by estimated EWMA mean (threshold at 25th/75th percentile, defining fast and slow contexts), compared with the static interval-response function. Shading indicates SEM across electrodes. Right: Intercepts of interval-response functions in fast and slow contexts. ***P < 0.001, two-tailed paired-sample t-test.

## DISCUSSION

Our study demonstrates an efficient neural code for time intervals in complex sound sequences such as speech. This neural code features (1) rapid adaptation to the mean and variance of the input time intervals and (2) stable compressive nonlinearity. The first feature highlights efficient neural coding based on the recent stimulus context, while the second feature suggests efficient neural encoding of positively skewed interval distributions that are commonly observed in natural sound. This neural code is observed in the temporal lobe around auditory cortex and its latency is around 100 ms, suggesting efficient time interval encoding in relatively early auditory activity.

### Context-dependent time encoding

Precise timing is essential when generating or perceiving complex motor sequences such as speech and music. In perception, timing cues can guide attentional orienting (Grabenhorst et al., 2019; Large & Jones, 1999; Nobre & van Ede, 2018; Schroeder & Lakatos, 2009) and the parsing of sequential structure (Bennett & Elfner, 2019; Frazier et al., 2006; Shukla et al., 2011). While the time intervals need to be precisely encoded, their range span more than an order of magnitude. We may encounter a syllable shorter than 50 ms or a musical note longer than 1000 ms (Fig. 1). Mapping such a wide range of intervals onto a limited neural dynamic range poses a challenge. This challenge can be alleviated when considering a specific context, e.g., speech from a particular speaker or a particular piece of music, and adapting the neural code based on the time interval distribution in a specific context (Fig. S1).

Here we demonstrate that the neural response flexibly adapts to the mean and variance of stimulus time intervals (Fig. 4). One previous study has shown that during single-interval timing, the population activity in primate prefrontal cortex adapts to the mean time interval in an experiment block (Meirhaeghe et al., 2021). Another study reveals that oscillatory neural activity from the human temporal lobe can track the rate of rhythmic speech and influence the perception of vowel length (Kösem et al., 2018). The current study, however, demonstrates sensory adaptation to both the mean and the variance of time intervals in connected sound sequences of different levels of rhythmicity. Critically, neural activity quickly adapts to the local context, as is revealed by the computational model and model-based MEG/iEEG analyses. Our model (Fig. 6) continuously infers the input distribution and modulates the interval-response function accordingly, offering an algorithmic implementation for rapid adaptation to variable contexts (Angeloni et al., 2023; Fairhall et al., 2001; Weber et al., 2019). This adaptive model also provides a plausible mechanism for rapid updating and error correction during sensorimotor synchronization (Large et al., 2023; Repp & Su, 2013; Vishne et al., 2021). Furthermore, it produces a negative serial dependency in subsequent interval estimates (Fig. S6 and Fig. S8), which is a hallmark of context-dependent timing behaviors such as duration adaptation (Corbett et al., 2021; Hayashi & Ivry, 2020; Heron et al., 2011; Johnston et al., 2006).

### Nonlinear time encoding

Time intervals in natural sound sequences are positively skewed (i.e., long-tailed) (Fig. 1) (Brudzynski, 2026; Greenberg et al., 2003; Hage et al., 2013). Efficient coding theory, however, posits that the neural response most efficiently encoding time intervals should be subject to an unskewed Gaussian distribution when the response power is limited (Cover & Thomas, 1999). Here, we demonstrate a compressive nonlinearity in the interval-response function (Fig. 5), which transforms a positively skewed input into a less skewed neural code. Therefore, from an evolutionary perspective, this compressive nonlinearity constitutes an efficient coding strategy for time intervals in natural sound (Sun et al., 2012). Furthermore, we show that the shape of this nonlinearity does not adapt to the skewness of the input distribution within the duration of our stimulus sequence, i.e., 20 s, reflecting a stable encoding strategy.

The compressive nonlinearity we observed is consistent with previous findings that suggest a roughly logarithmic scale for interval timing. For example, behaviorally, it has been shown that the subjective mean of time intervals is close to their geometric mean, which is equivalent to a direct average on a logarithmic scale (Kopec & Brody, 2010; Ren et al., 2020; Roberts, 2006). Furthermore, both human behavioral studies (Buhusi & Meck, 2005; Gibbon, 1977; Gibbon et al., 1984; Jazayeri & Shadlen, 2010) and studies recording rodent hippocampal activity (Cao et al., 2022; Mau et al., 2018) suggest that longer time intervals are encoded with less precision. Our study and these previous studies are both consistent with a compressive nonlinearity, but our study shows that the nonlinearity is better explained by a sigmoidal function rather than a logarithmic function.

### The efficient coding principle

The efficient neural principle has explained a wide range of sensory processing across species, for both visual features such as light intensity, contrast, and orientation (Brenner et al., 2000; Carandini & Heeger, 2012; Pitkow & Meister, 2012) and auditory features such as sound intensity, contrast, frequency, and spatial location (Dean et al., 2005; Herrmann et al., 2014; Rabinowitz et al., 2011; Willmore & King, 2023). Recent studies have extended it to more abstract cognitive domains, including value (Polanía et al., 2019; Schütt et al., 2024), number (Cheyette & Piantadosi, 2020; Prat-Carrabin & Woodford, 2026), and memory representations (Bays et al., 2024; Huang & Doeller, 2026). In line with these studies, our study suggests that the efficient coding principle applies to encoding time intervals, indicating a general computational mechanism by which efficient coding facilitates brain processing under tight resource constraints.

In natural sound, the mean and variance of time intervals are highly variable, while the right-skewness of time interval distributions is stable (Fig. 1). Consistent with the efficient neural encoding theory, neural adaptation to mean/variance and skewness shows distinct time scales: Rapid adaptation to mean and variance is reliably observed for each 20-second sequence, possible through the calculation of running average/variance (Fig. 6), whereas the nonlinearity is stable across stimulus distributions. A similar dissociation is found in auditory processing: the shape of the spectro-temporal receptive field is relatively stable, enabling extraction of evolutionarily relevant auditory features (Gervain & Geffen, 2019; Lewicki, 2002; Smith & Lewicki, 2006), whereas the response gain quickly adapts to the mean and variance of the sound intensity that supports robust listening under noisy backgrounds (Dean et al., 2005; Mesgarani et al., 2014; Rabinowitz et al., 2011). The difference in adaptation timescale may originate from the complexity and stability of the statistics (Fairhall et al., 2001; Lundstrom et al., 2008; Weber et al., 2019).

### Neural encoding of time intervals

The current study shows that the field potential in the auditory cortex encodes the duration of time intervals (Fig. 3). This result is consistent with previous findings that some auditory neurons show offset responses whose amplitude scales with the sound duration (Kopp-Scheinpflug et al., 2018; Qin et al., 2009). Similar offset responses that encode time intervals have also been observed in other modalities, including visual and tactile perception (Ofir & Landau, 2022; Wiener & Thompson, 2015). This neural code also aligns with the state-dependent model for time encoding (Goel & Buonomano, 2016; Motanis et al., 2018). This class of models posits that, when a sensory event occurs, it triggers a continuously evolving pattern of changes in the internal state of a neural network. Consequently, the amount of time that has passed since the event can be read out by examining where the neural network state sits along its trajectory. When processing a sound sequence, the amplitude of the evoked response to individual events is proportional to the interval preceding the event (A. L. Wang et al., 2008), and this effect can also contribute to the time-interval-sensitive response we observe. Critically, the neural encoding of time intervals in the current study adapts to the mean and variance of the temporal statistics, suggesting an active reconfiguration of the neural dynamics for efficient timing rather than a passive recovery process (Meirhaeghe et al., 2021).

Recent studies have also attributed speech timing to low-frequency neural oscillations in the human sensorimotor cortex that track the stimulus rhythm (Arnal & Giraud, 2012; Doelling et al., 2019; Poeppel & Assaneo, 2020; Zalta et al., 2024; Zoefel et al., 2018). The oscillation mechanism is proposed to underlie the processing of periodic stimuli (Lakatos et al., 2019; Obleser & Kayser, 2019), while it remains controversial whether such oscillations generalize to less rhythmic sequences (Oganian et al., 2023; Rimmele et al., 2018; Zou et al., 2021). The neural code observed here applies to both aperiodic (e.g., gamma distribution with α = 1) and periodic (e.g., gamma distribution with α = 16) stimuli, while the oscillatory code may be specialized to more periodic stimuli.

Taken together, our findings suggest that the human brain utilizes the efficient coding principle to encode time intervals in sound sequences. The neural code adapts to the statistical properties of natural sounds, and the timescale of adaptation is aligned with the timescale of variability of statistical properties. These results reveal a neural mechanism that supports precise and flexible time encoding for sound sequences.

## METHODS

### Temporal statistics of time intervals in natural sounds

Six speech corpora were included in the analysis, i.e., TIMIT (Garofolo et al., 1993), AISHELL-1 (Bu et al., 2017), GigaSpeech Talk/Audiobook (Chen et al., 2021), and WenetSpeech Talk/Audiobook (B. Zhang et al., 2022), covering two languages (i.e., English and Chinese) and three speaking styles (reading sentences, reading audiobooks, and spontaneous speech). Boundaries between phonemes were extracted based on the audio and transcription using the Montreal Forced Aligner (MFA) (McAuliffe et al., 2017; Y. Zhang et al., 2023). The MFA directly provided the syllable boundaries for Chinese. For English, the syllable boundaries were determined by grouping phonemes into syllables based on a dictionary, i.e., the Unisyn Lexicon (Fitt, 2001). We then calculated the mean, variance, and skewness of the syllable durations for each speaker. We also characterized other natural sound sequences, including human songs in nine languages (Y. Zhang et al., 2024), the birdcall of four species (Cohen, 2022; Hedley, 2016; Lachlan et al., 2018; Nicholson et al., 2017), and human footsteps (Larracy et al., 2025). We extracted the skewness of each song in human song dataset, each bird in birdcall dataset, and each individual in footstep dataset.

### Participants

Sixty native Chinese listeners (18 to 25 years old, mean age, 21.8 years) participated in the MEG Experiment 1 (N = 20, 10 males), Experiment 2 (N = 20, 13 males), and Experiment 3 (N = 20, 9 males). All participants were right-handed (Oldfield, 1971) with no self-reported hearing loss or neurological disorders. All participants gave written informed consent before enrollment and received monetary compensation. The experimental procedures were approved by the Ethics Committee of the College of Biomedical Engineering and Instrument, Zhejiang University (No. 2022–001).

### MEG experimental stimuli

The auditory stimuli in MEG experiments were constructed in three steps (Fig. 2e). First, a sequence of time intervals was either extracted from natural speech recordings or sampled from a given statistical distribution (see *Generation of time intervals*). Second, the time interval sequences were temporally scaled into two mean syllable rates (the average number of syllables per second). Third, each interval was filled with a syllable /sa/ (see *Generation of syllable sequence stimuli*). All stimuli were delivered binaurally via air-tube earphones at a sampling rate of 44,100 Hz. In the following, we first described how the time intervals were generated and then how a sound sequence was generated based on the time intervals.

#### Generation of time intervals

##### Experiment 1

We tested sequences constructed based on syllable onset/offset times in natural speech. Syllable onsets and offsets were first derived from four speech corpora: AISHELL-1 (Chinese, read sentences), WenetSpeech-talk (Chinese, spontaneous speech), TIMIT (English, read sentences), and GigaSpeech-talk (English, spontaneous speech). For each corpus, we selected four utterances, each containing more than 100 syllables, with the constraint that the mean syllable rate was between 3 and 5 Hz. For AISHELL-1 and TIMIT, sentences read by the same speaker were concatenated to form a longer utterance. The extracted time intervals were linearly scaled to create two mean syllable rates, i.e., 3 Hz (mean duration: 333 ms) and 5 Hz (mean duration: 200 ms). For each syllable rate, we kept a minimal integer number of syllables so that the sequence was at least 20 seconds long (approximately 60 syllables for 3 Hz and 100 syllables for 5 Hz).

##### Experiment 2

To separately manipulate the mean and variance of time intervals, we sampled time intervals from the Gamma distributions with five shape parameters, i.e., α = 1, 2, 4, 8, and 16. In each stimulus, all time intervals were independently generated based on a Gamma distribution of a fixed shape parameter with a mean syllable rate of 5 Hz. The sampled time interval sequences were linearly scaled to create a corresponding sequence with a mean syllable rate of 3 Hz. In gamma distributions, when the mean rate was fixed, a larger *α* led to a distribution with smaller variance. Sampled time intervals shorter than 50 ms were set to 50 ms to ensure that the syllable generated would be clearly audible. For each combination of syllable rate and shape parameter, two independent sequences were generated, and we kept a minimal integer number of syllables so that the sequence was at least 20 seconds long (approximately 60 syllables for 3 Hz and 100 syllables for 5 Hz).

##### Experiment 3

To manipulate the skewness of the duration distribution, we employed two distribution types: uniform and skew-normal distributions. The mean syllable rates were set to 3.3 Hz and 4.5 Hz, which were closer than those used in Experiments 1 and 2, thereby increasing the difficulty of the fast/slow discrimination task and maintaining participant engagement. For the uniform distribution (skewness = 0), duration ranges were set from 140 to 470 ms for 3.3 Hz and from 90 to 350 ms for 4.5 Hz. For the skew-normal distribution, we used the skewnorm function in Scipy (Virtanen, 2020) with the parameters (a = −100, loc = 0.43, scale = 0.17) for 3.3 Hz and (a = −100, loc = 0.33, scale = 0.15) for 4.5 Hz, achieving a skewness of −0.6 while maintaining variance similar to the uniform distribution. Sampled durations shorter than 50 ms were thresholded at 50 ms to ensure reliable syllable discriminability. For each combination of syllable rate and distribution type, five independent sequences were generated, and we kept a minimal integer number of syllables so that the sequence was at least 20 seconds (approximately 66 syllables for 3.3 Hz and 90 syllables for 4.5 Hz).

#### Generation of syllable sequence stimuli

To generate the syllable /sa/ at different durations, we first recorded a short and a long exemplar of /sa/ and edited them in Praat (Boersma, 2001). The fricative /s/ segment was extracted from the short exemplar and scaled to 30 ms. The vowel /a:/ segment was extracted from the long exemplar and scaled to various durations from 20 ms to 1500 ms. The scaling of duration was achieved using Praat’s built-in Compressor function, which preserved the pitch and spectral characteristics during temporal manipulation. The consonant and vowel segments were concatenated to create /sa/ syllables of the target duration. Syllables were then concatenated sequentially according to the scaled duration sequences to generate continuous auditory stimuli.

### MEG experimental procedure

The experimental paradigm was implemented using Psychophysics Toolbox-3 for MATLAB (MathWorks). In all three experiments, participants performed a speed discrimination task: on each trial, they listened to a syllable sequence and judged whether its mean rate was “fast” or “slow” via button press. The inter-trial interval was randomly sampled between 1 s and 1.5 s. The experimental setup for each experiment will be detailed in the following. Before the main experiment, participants completed a training phase using stimuli from the same conditions in the formal session but with different samples. Participants advanced to the formal session after achieving at least 80% accuracy.

#### Experiment 1

Experiment 1 comprised 128 trials from 8 conditions (2 rates × 4 corpora). For each corpus, the four syllable sequences were divided into two pairs (1-2 and 3-4) and counterbalanced across participants: for half of the participants, sequences 1 and 2 were assigned to 3 Hz conditions and sequences 3 and 4 to 5 Hz conditions; for the other half, this assignment was reversed. As a result, each participant heard 16 stimuli (4 corpora × 2 sequences × 2 rates), and each stimulus was repeated 8 times (16 stimuli × 8 repetitions = 128 trials). All trials were randomly intermixed during presentation. Participants took a short break after every 32 trials.

#### Experiment 2

Experiment 2 comprised 120 trials from 10 conditions (2 rates × 5 gamma distributions). Each condition contained 2 independent stimulus sequences, and each stimulus was repeated 6 times (10 conditions × 2 sequences × 6 repetitions = 120 trials). All trials were randomly intermixed during presentation. Participants took a short break after every 30 trials.

#### Experiment 3

Experiment 3 comprised 120 trials from 4 conditions (2 rates × 2 distribution types). Each condition contained 5 independent stimulus sequences, and each stimulus was presented 6 times (4 conditions × 5 sequences × 6 repetitions = 120 trials). All trials were randomly intermixed during presentation. Participants took a short break after every 30 trials.

### MEG acquisition and pre-processing

Neuromagnetic responses were recorded using a 306-sensor whole-head MEG system (Elekta-Neuromag, Helsinki, Finland) at Shenzhen University, sampled at 1 kHz. The system had 102 magnetometers and 204 planar gradiometers. Four MEG-compatible electrodes were used to record EOG at 1 kHz. Four head position indicator (HPI) coils were used to measure the head position inside MEG. The positions of three anatomical landmarks (nasion, left, and right pre-auricular points), the four HPI coils, and at least 250 points on the scalp were also digitized before the experiment.

The recorded MEG data were preprocessed offline utilizing MNE-Python tools (Gramfort et al., 2013). Initially, the data underwent de-noising and motion correction using the Maxfilter Temporal Signal Space Separation (tSSS) method. To remove ocular artifacts in MEG, the horizontal and vertical EOG were regressed out from the recordings using the least-squares method. Subsequently, the data were band-pass filtered between 0.3 and 40 Hz using a linear-phase finite impulse response (FIR) filter (-6dB attenuation at the cut-off frequencies, 10 s Hamming window), and downsampled to a sampling rate of 100 Hz. Only gradiometers were included in the subsequent analysis.

### iEEG dataset

#### English natural speech

For English natural speech listening, we used the “Podcast” dataset (Zada et al., 2025). This dataset utilized electrocorticography (ECoG) to record neural activity from nine participants while they listened to a 30-minute narrative podcast (“So a Monkey and a Horse Walk Into a Bar, Act One: Monkey in the Middle”). The audio story contained 5,136 words and 7,228 syllables, with the word and syllable boundaries manually annotated. ECoG signals were recorded from a total of 1,268 electrodes (1,057 over the left hemisphere, 211 over the right) at a sampling rate of 512 Hz. Low-frequency signal (band-pass filtered between 0.1–40 Hz) was extracted and downsampled to 100 Hz for analysis.

#### Chinese natural speech

For Chinese natural speech listening, we used the “The straw house” stereotactic electroencephalography (sEEG) dataset presented in our previous work (Y. Wang et al., 2025). This dataset utilized sEEG to record neural activity from 19 native Chinese speakers while they listened to an 80-minute continuous audiobook narrative (“The Straw House”). The audio story contained 14,528 Chinese characters (i.e., syllables) and was manually annotated with precise onset times. sEEG signals were recorded from a total of 2,606 valid electrodes (1,453 over the left hemisphere, 1,153 over the right) at a sampling rate of 2,048 Hz. Low-frequency signal (band-pass filtered between 0.1–40 Hz) was extracted and downsampled to 100 Hz for analysis.

### Modeling interval-response function

#### Time-resolved correlation

We used time intervals to predict neural data for each channel in MEG and each electrode in iEEG. To isolate interval-encoding neural activity, the neural response to basic acoustic features was regressed out first using the temporal response function (TRF) (Ding & Simon, 2012) before further analysis (see Supplementary Information). Neural activity was then aligned based on the offset of each syllable, and averaged using a 30-ms sliding window across 100 time lags, ranging from −500 ms to +500 ms in 10-ms steps. Time-resolved correlation was independently calculated for each stimulus condition and each time lag. Considering that the brain may encode time intervals in a nonlinear manner, we tested three regression functions, including a linear function, a logarithmic function, and a sigmoidal function. These functions constrained how the brain encodes the time intervals. The logarithmic function applied a logarithmic transformation log(*x*+*a*) to the intervals before linear regression (*a* as a free parameter to control the function shape). The sigmoidal function was defined by four parameters controlling the lower bound, upper bound, slope, and inflection point of the function (*l, u, a, x_0_*):

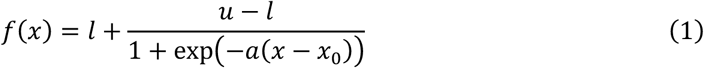

The three functions were separately fitted to predict neural responses to time intervals. We used 4-fold cross-validation to assess the performance of models. We evaluated out-of-sample prediction by computing the Pearson correlation between the predicted and the actual signal for each held-out test set. The best-performing model, as determined by the cross-validated Pearson correlation, was used for all subsequent adaptation analyses.

#### Interval-response function

We integrated data from whole-brain channels to obtain one interval-response function for each participant that was used in subsequent adaptation analysis. To avoid excessive data selection, we calculated a unified peak time lag over all conditions for each participant. All peak time lags were distributed between 100 ms and 120 ms. We then identified the five channels with the highest encoding correlations at this peak time lag as regions of interest (ROIs) for each participant. For these five channels, we extracted the neural responses to each time interval at the time lag. We corrected the polarity of responses to ensure a positive correlation between neural activity and time intervals for each channel, then averaged across all channels to obtain each participant’s integrated neural response to time intervals. Finally, we fitted a sigmoid function to derive the parameters of the integrated interval-response function. The use of five channels as ROIs balanced noise reduction and signal strength and optimized the correlation between time intervals and neural responses. However, the adaptation analysis in the following was robust to the number of ROIs: using anywhere from 1 to over 10 channels did not affect the main conclusions of the study.

### Analyses of interval-response function

#### Analysis of adaptation to mean and variance

For each participant, we first computed a global mean response level r_mean_ by averaging neural responses to all syllables across all conditions. For each condition, we then defined the intercept *b* as the time interval whose neural response equals r_mean_, i.e., b = *f* ^-1^(r_mean_), where *f* (·) was the interval-response function fitted under that condition. The intercept indexed the horizontal location of the response function on the stimulus axis. For each mean syllable rate, we averaged the intercepts of all conditions within this rate to obtain the mean intercept. Regarding variance, we tested the local slope at the intercept, i.e., the derivative of the interval-response function at the intercept: *k_b_* = *f*’(*b*). We then averaged the slopes across all conditions within each rate to obtain the mean slopes. In Experiment 2, we additionally analyzed the Pearson’s correlation between the slopes of interval-response functions and the reciprocal of the standard deviation of the gamma distributions within each rate.

#### Analysis of adaptation to skewness

Under the efficient coding hypothesis, successful neural encoding should make the output distribution’s skewness closer to zero compared to the input distribution. To test this, we directly compared the distributions of neural activity and input stimuli. A methodological challenge was that neural noise can artifactually reduce skewness, confounding the effect of the nonlinear transformation. To address this, we equated noise levels between stimulus and response by adding calibrated noise to the stimulus distribution. For each participant and condition, we computed inter-trial correlation (ITC) between neural activity across different trials as a measure of signal-to-noise ratio. We then added Gaussian noise to the stimulus to match its ITC with that of the neural response. Specifically, we added i.i.d. Gaussian noise *N*(0, *σ*²) to the stimulus duration, and identified σ via binary search such that the ITC computed on the noised stimulus (averaged across 100 samples) matched the neural ITC within 0.00001. After that, we calculated the skewness of both neural activity and the noised stimulus (averaged across 100 samples). This model-free approach avoided biases introduced by designated encoding models. Lastly, we examined the shape of the interval-response function as a model-based validation. We calculated the second derivative of the interval-response function at the intercept, *f*’’(b), to quantify its influence on the skewness. A negative second derivative indicated a nonlinearity that compressed the long intervals and decreased the skewness.

### Dynamic adaptation model

#### Model description

We constructed a dynamic adaptation model to investigate the adaptation process underlying neural time interval encoding. The model comprised two components: an adaptive encoder with adjustable parameters and an observer that estimates the mean and standard deviation of the input distribution.

For the encoder, we assumed that it performed an intercept shift and gain control based on the estimated mean and standard deviation. We implemented the adaptive encoder as a linear-nonlinear model: *r*_t_ = *f*(*k*(*s_t_*-*b*)). The *k*(*s_t_*-*b*) represents the linear scaling, and *f*(·) is the sigmoid function defined above. The slope *k* and intercept *b* are scaled with the estimated standard deviation and mean:

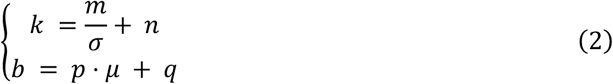

For the observer, we employed an exponentially weighted moving average (EWMA) model to estimate both μ and σ from the stimulus history. The exponential-averaging model computes smoothed estimates of μ and σ by taking a weighted average of previously perceived time intervals, giving more weight to recently experienced intervals:

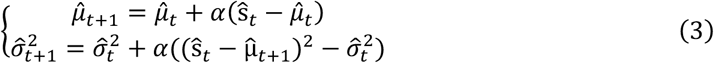

where *α* is the learning rate governing the weight of new observations. In MEG experiments, the initial estimates μ_0_ and σ_0_ were set to the global mean and standard deviation of all stimulus intervals across conditions. In iEEG datasets, the initial estimates *μ*_0_ and *σ*_0_ were set to the mean and standard deviation of the entire speech stimuli. This model required minimal parameters and memory storage, enabling highly efficient estimation of stimulus statistics. Moreover, this model has been shown to provide optimal fits to human parameter estimation behavior in numerous previous studies (Landy et al., 2012; Norton et al., 2017, 2019).

To fit the dynamic adaptation model to neural data, we adopted a two-stage iterative fitting procedure. For a fixed observer learning rate α, we used least-squares optimization to identify the optimal combination of scaling parameters (*m, n, p, q*) and nonlinear activation function parameters (*l, u, a, x*_0_). The optimizer employed the Trust Region Reflective algorithm (Byrd et al., 1988). Initial values for (*l, u, a, x0*) were obtained by fitting a single sigmoid to the pooled data of all conditions; (*m, n, p, q*) were initialized using 10 random initial points and then optimized to get the best parameters. The bounds of (*l, u, a, x0, m, n, p, q*) were between -100 and 100, and the bounds of (a, m, p) were between 0 and 100 to ensure the direction of dynamic adaptation in line with efficient coding. Subsequently, we used Bayesian optimization (60 iterations) to iteratively optimize the learning rate *α* ∊ (0, 1) that maximizes the correlation between predicted and actual neural response. Model performance was evaluated using the cross-validation correlation: in MEG Experiments 1 and 2, where each condition contained 2 independent stimulus sequences, we used 2-fold cross-validation, with one sequence per condition serving as training and the other as test in each fold; in Experiment 3, where each condition contained 5 independent sequences, we used 5-fold cross-validation. In the iEEG dataset, we first divided the speech into 5 continuous pieces and then used the 5-fold cross validation. Model performance was quantified as the Pearson correlation between predicted and observed neural activity.

#### Alternative Models

We compared the dynamic adaptation model against alternative time interval encoding models. In the static model, a single interval-response function was fitted across all conditions to the neural response, assuming no context-dependent adaptation. In the oracle model, separate interval-response functions were fitted to the neural response for each condition independently, representing a condition-specific encoding with perfect knowledge of the current trial’s statistics. We also evaluated several reduced versions of the dynamic adaptation model (Fig. S4). Among these reduced models, EWMA_mean_ adapted to only the mean estimate (intercept change) while keeping the slope fixed, and EWMA_std_ adapted to only the variance estimate (slope change) while keeping the intercept fixed. Another control model, EWMA_shuffle_, estimated the mean and variance from temporally shuffled stimulus contexts. The shuffle was implemented within each trial, which disrupts the temporal order while preserving the overall statistics. To avoid overfitting and facilitate model comparison, all model fitting was performed using the correlation between predicted and actual neural responses with the same cross-validation in the EWMA model.

#### Context sensitivity analysis

We tested whether the neural encoding adapts to the within-trial fluctuations of estimated statistics. Based on the EWMA-estimated mean at each step, we identified intervals in two types of local contexts within each trial: intervals in which the estimated mean was below the 25^th^ percentile of the estimated mean distribution (fast context) and intervals in which the estimated mean was above the 75^th^ percentile of the estimated mean distribution (slow context). For each participant and each condition, we refitted the interval-response function separately in the fast context and slow context and extracted the intercept *b*. We define the difference between intercepts in fast and slow contexts (averaged across conditions) as a measure of neural sensitivity to local context. In iEEG dataset, the entire speech was viewed as a trial to identify intervals in fast and slow contexts.

### Statistical analysis

#### Analysis of time-resolved correlation

In MEG experiments, we used a permutation t-test with 5,000 permutations to assess whether the peak correlation between syllable duration and neural responses was significantly higher than 0 for each channel. We compared the peak encoding correlations of different encoding models using paired t-tests. We used a one-dimensional cluster-based permutation test (5,000 permutations) to determine the timepoints with significant neural encoding of time intervals. The correlation used for model comparison and detecting significant timepoints was the averaged correlation over the five top channels for each participant (see *Interval-response function* for the selection of channels). In iEEG datasets, we identified significant electrodes by first thresholding the encoding correlation based on a permutation-based test. We permuted the mapping between input intervals and neural responses 10,000 times to obtain a null distribution of peak correlation and a P value was computed as the percentile of the actual peak corelation from the null distribution. Electrodes with P values less than 1×10^-4^ were chosen with mean correlation threshold = 0.057. Considering the electrodes with adaptive neural coding of time intervals should exhibit precise encoding power, we further used a stricter correlation threshold of 0.10 to select significant electrodes.

#### Analysis of interval-response function

To test the adaptation of the interval-response function to the mean and variance of the interval distribution, we used paired t-tests to compare the differences in intercept and slope between the two rates in each experiment. Additionally, in Experiment 2, we used Pearson’s correlation to test whether the slope of the interval-response function was correlated with the reciprocal of the interval standard deviation. To test the adaptation of the interval-response function to the skewness of the interval distribution, we used paired t-tests to compare the differences in the skewness and absolute skewness of intervals and neural responses for each condition. In addition, we compared the second derivatives of interval-response functions at the intercept versus 0 using a t-test for each condition.

#### Analysis of dynamic adaptation model

We compared the encoding correlations of EWMA models with alternative models using paired t-tests, with FDR for multiple comparison correction. In the context-sensitivity analysis, we used a paired t-test to assess whether the intercept differed significantly between the fast and slow contexts. In MEG experiments, the intercepts of fast and slow contexts were averaged across all conditions.

When multiple comparisons were performed in the statistical analyses above, P values were adjusted using the FDR correction (Benjamini & Hochberg, 1995).

## Data and code availability

All data and code in this study will be available upon publication.

## Acknowledgments

This work was supported by the National Natural Science Foundation of China (No. 32595493). We thank the members of the Ding Lab for thoughtful discussions and feedback. We thank M. Zhang, M. Fang, and the Center for MEG at Shenzhen University for assistance with data collection.

## Supplementary Information

### S1 Simulation of the efficient encoding strategies for time intervals in natural speech

We simulated four strategies to encode time intervals: static encoding for all time intervals, adaptation to the mean of each speaker, adaptation to the mean and variance of each speaker, and the maximum entropy strategy that outputs Gaussian distributions for each speaker. These adaptation strategies show incremental entropy of the simulated neural code, demonstrating the coding efficiency.

**Figure S1:**
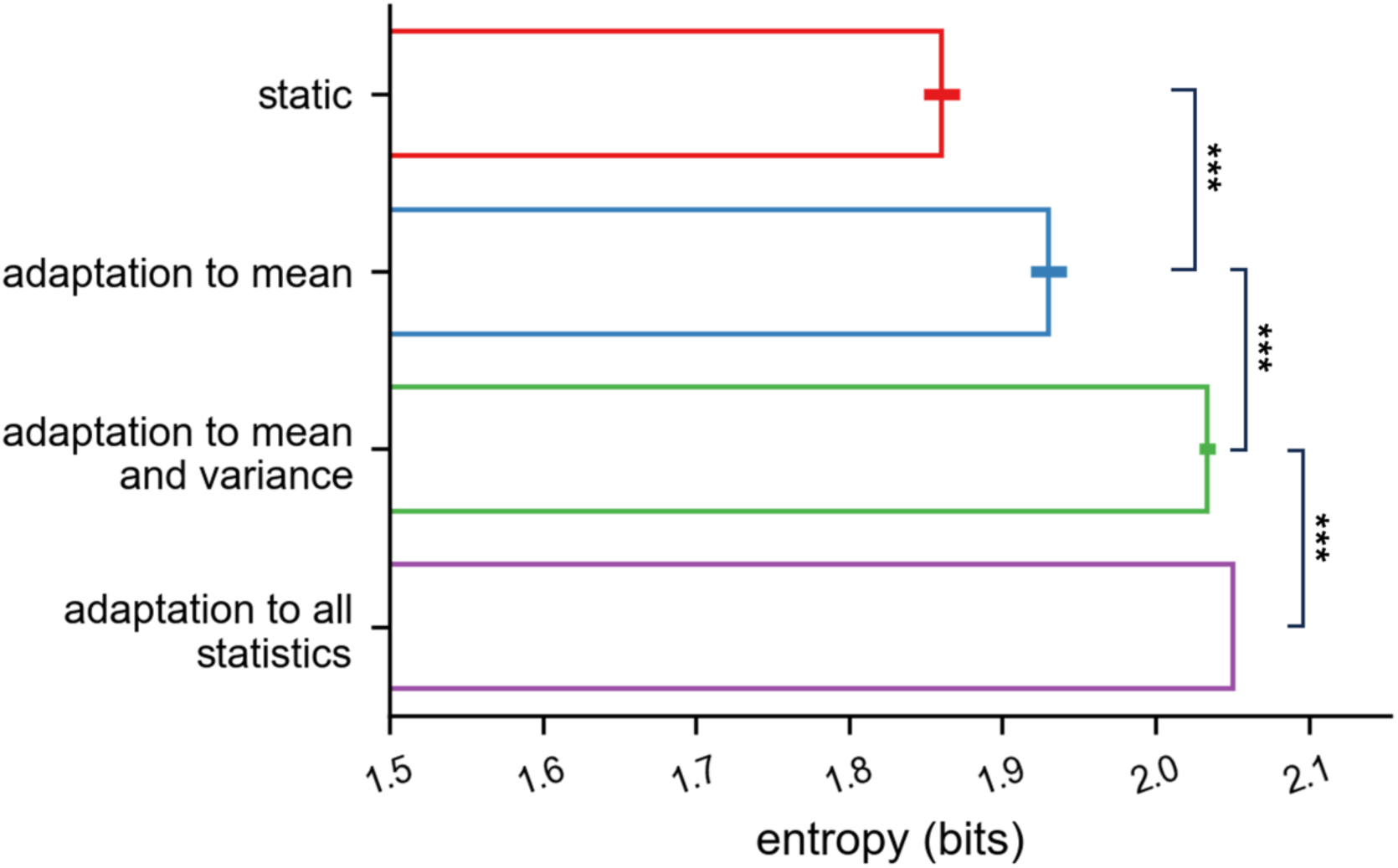
Simulation of the efficient encoding strategies for time intervals in natural speech. Error bars indicate SEM of speakers. ***P < 0.001, two-sided t-test with FDR correction.

### S2 Neural encoding of time intervals in Experiments 2 and 3

**Figure S2:**
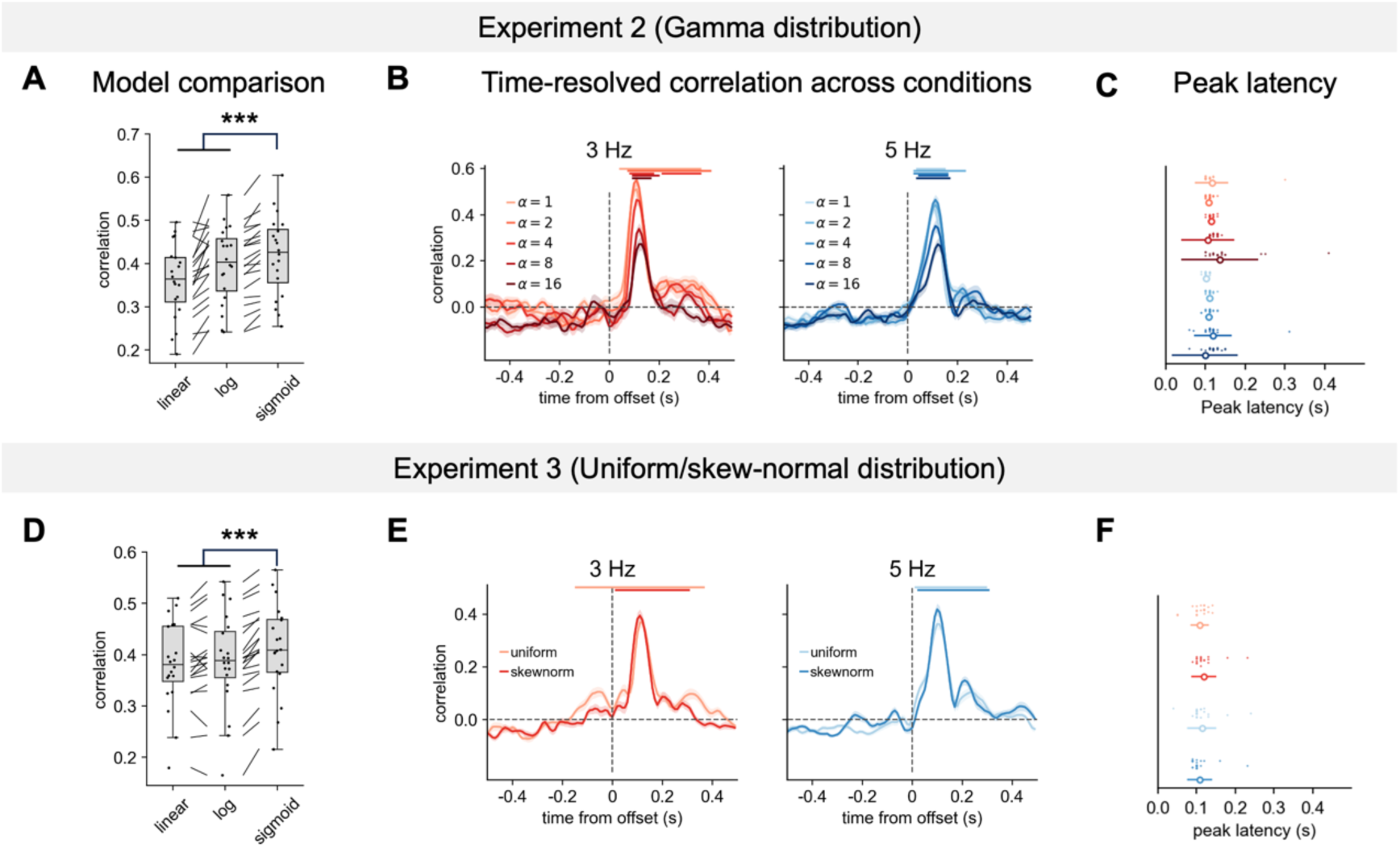
Neural encoding of time intervals in Experiments 2 and 3. (A) Peak correlation for three curve fits of interval-response function in Experiment 2. Whisker plots show the distribution of correlation values (center line, median; box, 25th–75th percentile; whiskers, ±1.5 × interquartile range; dots, individual participants). The sigmoidal model outperforms linear and logarithmic models (***P < 0.001, two-sided t-test with FDR correction). (B) Time-resolved correlation between MEG response and syllable duration for 3 Hz (left) and 5 Hz (right) conditions. Shading indicates SEM Horizontal bars denote significant encoding (cluster-based permutation test, P < 0.05; cluster-forming using point-wise t-test P < 0.01 with 5,000 permutations). (C) Peak latency of encoding correlation for all conditions. Error bars, s.d. across participants. (D-F) Same as (A-C) for Experiment 3.

### S3 Topography of adaptation to duration mean and variance

**Figure S3:**
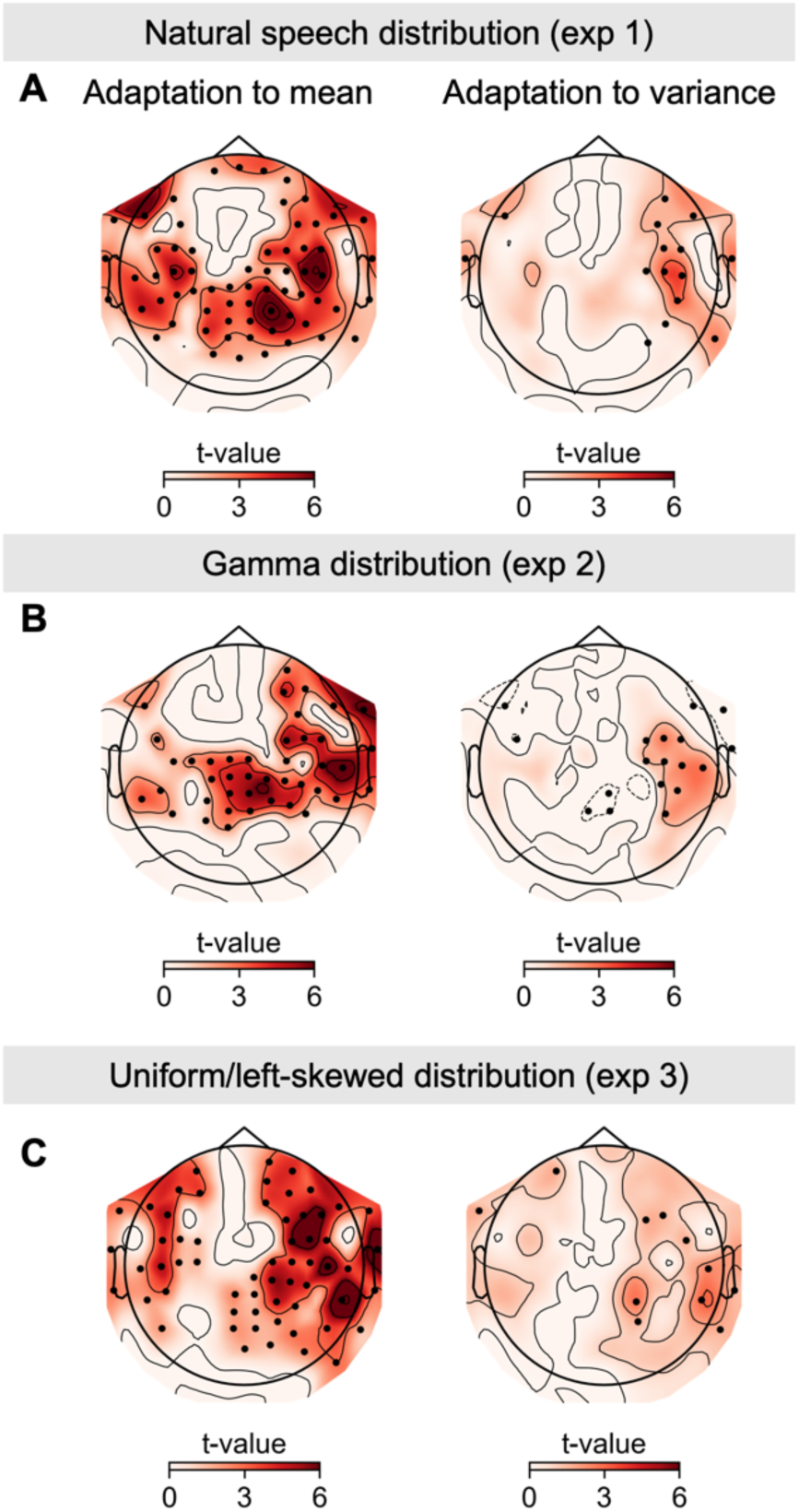
Topography of adaptation to mean and variance of time interval distribution. (A) Adaptation of the intercept (left) and slope (right) of interval-response functions to the mean and variance of time intervals for 3 Hz and 5 Hz conditions in Experiment 1. Dotted channels show significant adaptation, P < 0.05, permutation t-test with 5,000 permutations, FDR-corrected. (B-C) Same as (A) for Experiments 2 and 3.

### S4 Extended Model Comparison in Experiments 1, 2, and 3

Except for the static and oracle model, models that estimated only the mean (EWMA_mean_) or only the variance (EWMA_std_) performed significantly worse than the full model, as did a control model that estimated statistics from shuffled contexts (EWMA_shuffle_). These results confirm the continuous estimation of both mean and variance in the human brain.

**Figure S4:**
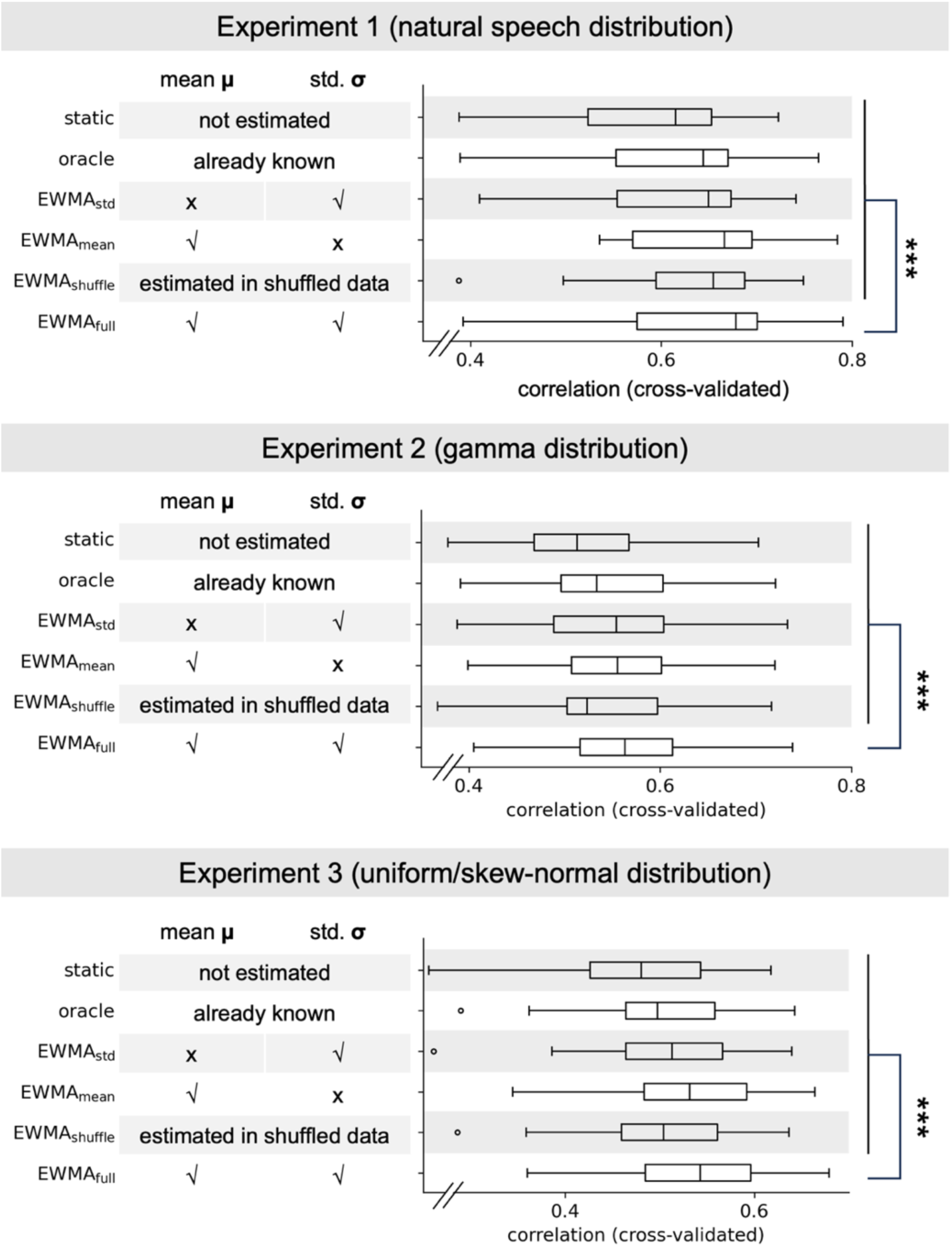
Extended model comparison. (A) Six models with different context estimation strategies (shown in the table) were fitted to the data. Box plots show cross-validated correlations (center line, median; box, 25th–75th percentile; whiskers, ±1.5 × interquartile range). The EWMA model outperforms all alternatives (***P < 0.001, two-sided t-test with FDR correction). (B-C) Same as (A) for Experiments 2 and 3.

### S5 Within-trial Dynamics in Experiments 2 and 3

**Figure S5:**
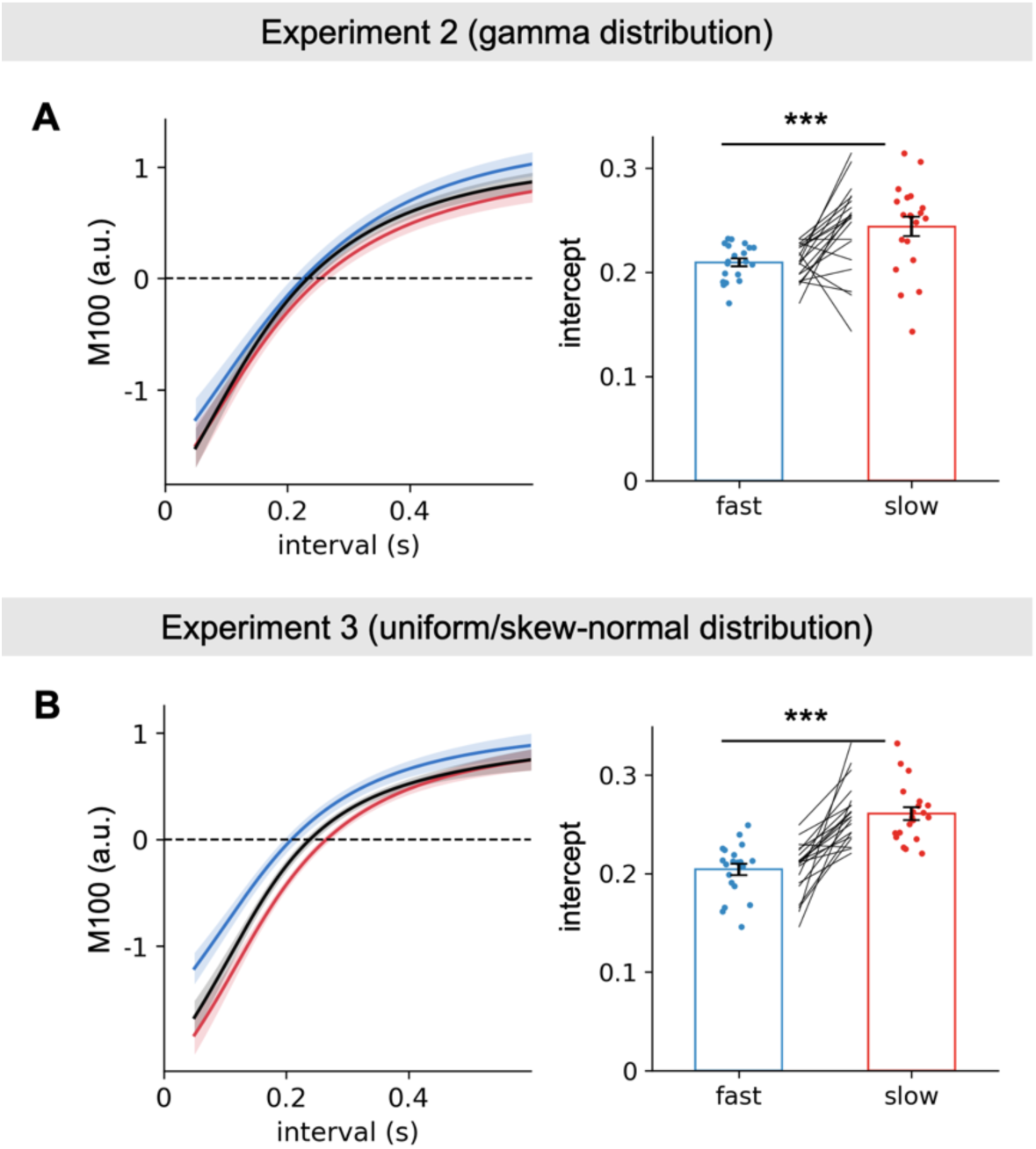
(A) Context sensitivity across participants in Experiment 2. Left: Interval-response functions fitted separately for fast and slow contexts, averaged across conditions and participants; shading indicates SEM of participants. Right: Intercepts of the interval-response functions for fast and slow contexts. ***P < 0.001, two-tailed paired-sample t-test. (B) Same as (A) for Experiment 3.

### S6 Serial dependency analysis in Experiments 1, 2, and 3

The serial dependency analysis (see *S9* for detailed description) showed that neural responses were significantly modulated by earlier intervals, with predominantly negative weights (Fig. S6 A-C; two-sided t-test, FDR-corrected P < 0.001). The EWMA model successfully reproduced these serial dependency patterns (Fig. S6 D-F).

**Figure S6:**
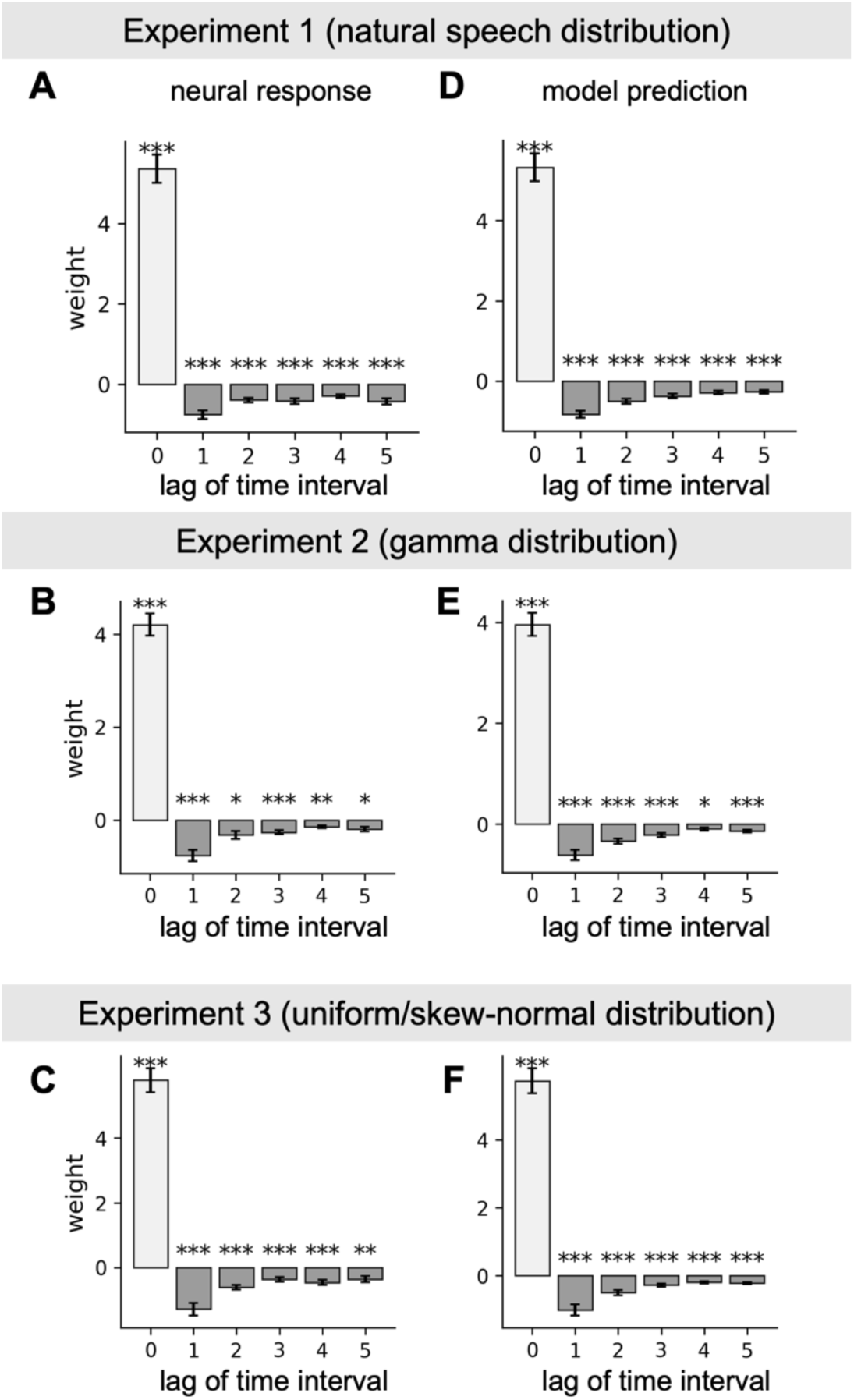
(A-C) Serial dependency of neural response on the current interval (lag 0) and preceding intervals (lag > 0) in Experiments 1, 2, and 3. (D-F) Serial dependency of the EWMA model prediction on the current interval (lag 0) and preceding intervals (lag > 0) in Experiments 1, 2, and 3. *P < 0.05, **P < 0.01, **P < 0.001, two-tailed paired-sample t-test with FDR correction.

### S7 Efficient Coding of Time Intervals in the Chinese Natural Speech Dataset

**Figure S7:**
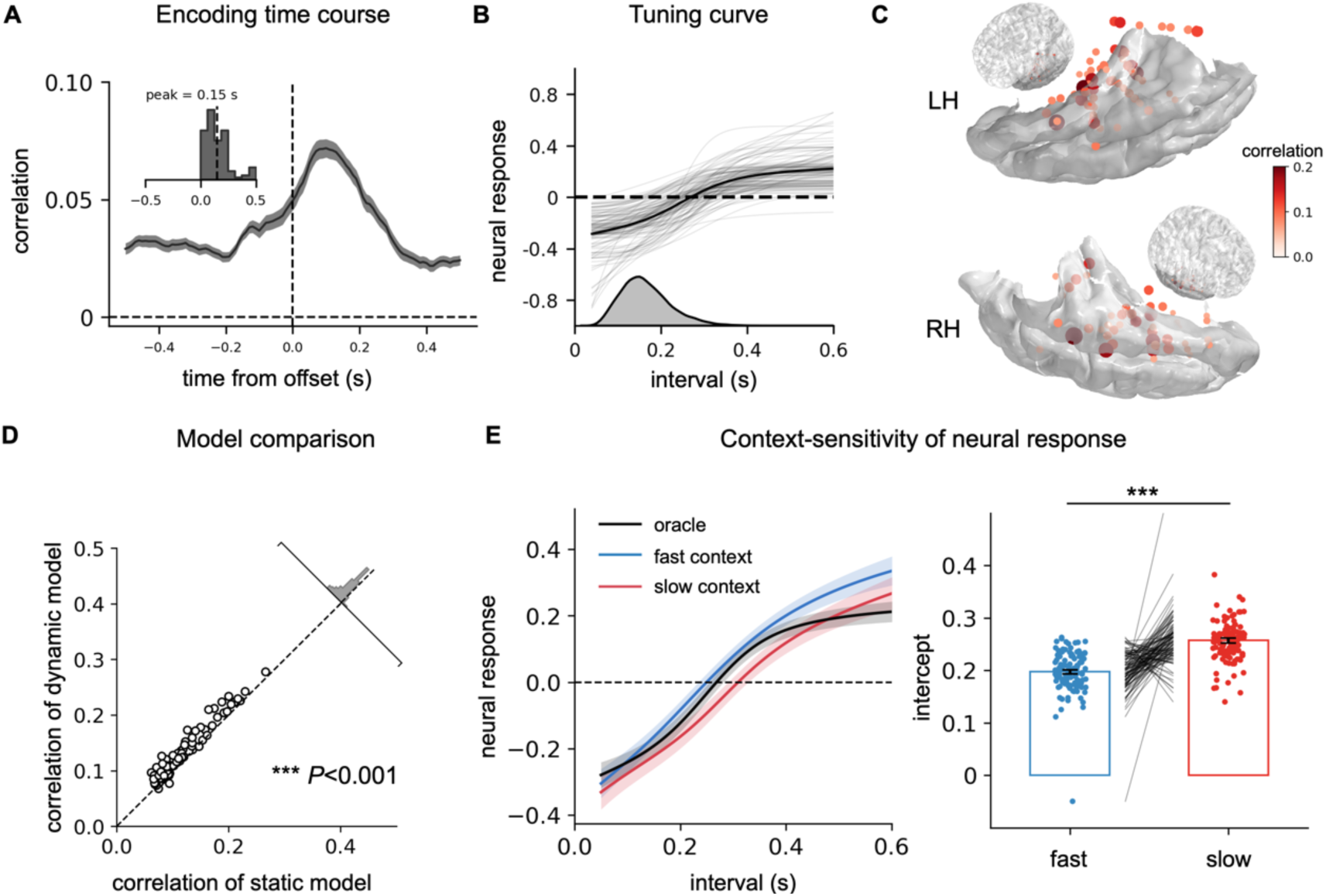
Efficient neural coding of time intervals during Chinese natural speech comprehension. (A) Time course of encoding correlation for significant electrodes (N = 108). Inset: distribution of peak encoding latencies across electrodes. (B) Interval-response function for significant electrodes. The histogram below shows the interval distribution. (C) Spatial distribution of encoding strength across electrodes. (D) Model comparison between the dynamic adaptation model and the static sigmoidal model. ***P < 0.001, two-tailed paired-sample t-test. (E) Context-dependent neural responses across electrodes. Left: Interval-response functions for intervals classified by estimated EWMA mean (threshold at 25th/75th percentile, defining fast and slow contexts), compared with the static interval-response function. Shading indicates SEM Right: Intercepts of interval-response functions in fast and slow contexts. ***P < 0.001, two-tailed paired-sample t-test.

### S8. Serial dependency analysis in two iEEG datasets

The serial dependence of neural activity on preceding intervals also showed significantly negative weights that decayed gradually (Fig. S8A-B); two-sided t-test, FDR-corrected P < 0.001). Furthermore, individual electrodes exhibited diverse learning rates of EWMA (Fig. S8C-D), with higher learning rates predicting stronger weighting of recent history, corresponding to faster adaptation (Fig. S8E-F).

**Figure S8:**
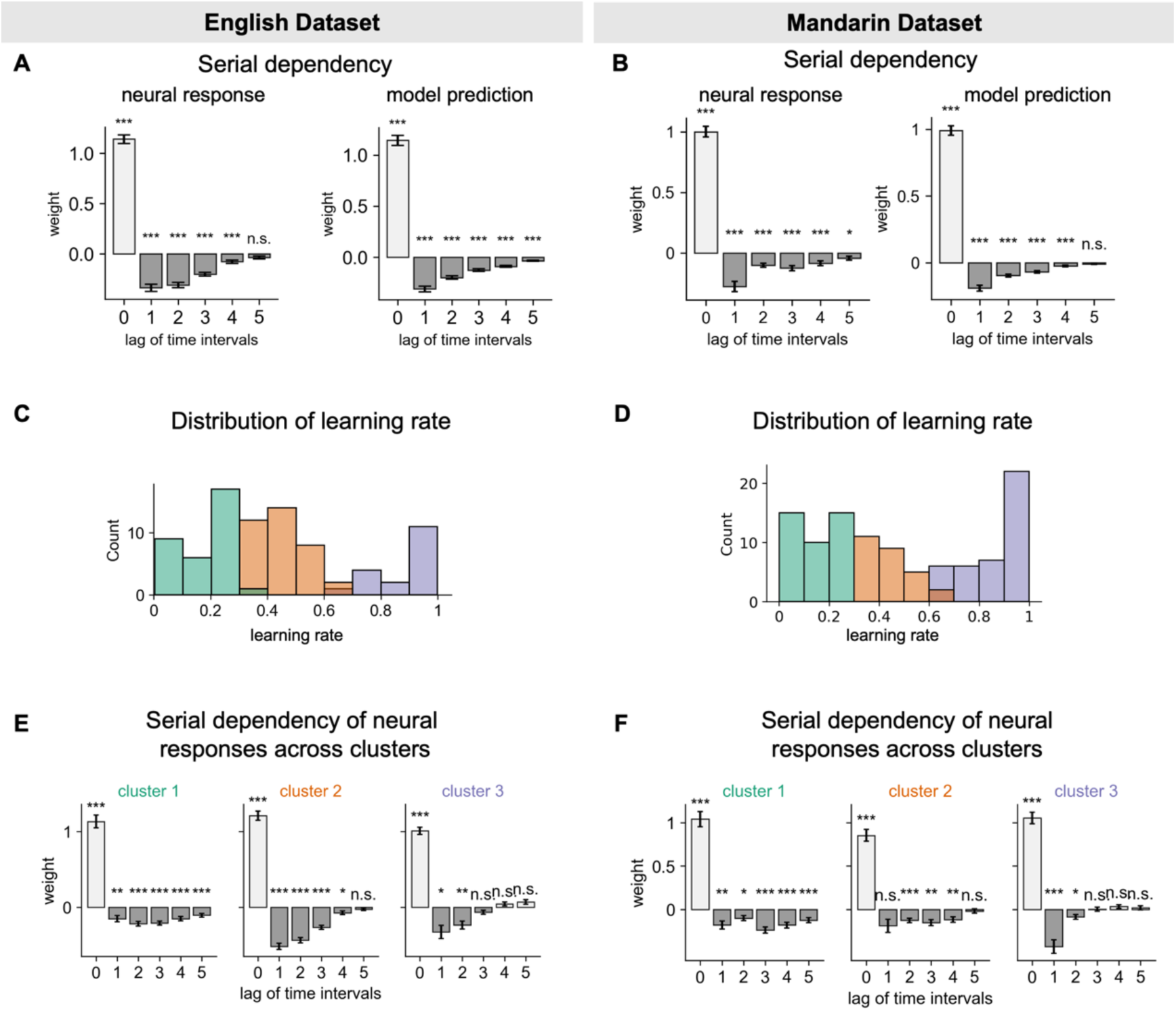
(A-B) Serial dependency of neural response and model prediction on the current interval (lag 0) and preceding intervals (lag > 0) in English and Chinese Dataset. (C-D) Clustering of the EWMA model learning rate fitted in significant electrodes using k-means with k = 3. (E-F) Serial dependency of neural response on the current interval and preceding intervals across three clusters. *P < 0.05, **P < 0.01, **P < 0.001, two-tailed paired-sample t-test with FDR correction.

### S9 Extended methods

#### Dissociating acoustic confounding through Temporal response function (TRF)

We used TRF to regress out the acoustic response from the neural response (Ding & Simon, 2012). The TRF was employed to model the time-domain relationship between the sound envelope and the neural response. The model assumes that the neural response to the sound envelope can be described as:

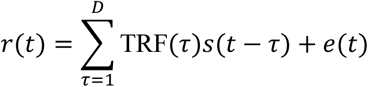

where *r*(*t*), *s*(*t*), and *e*(*t*) denote the MEG response, the envelope of speech, and the residual error, respectively. The envelope was extracted by an auditory model that filtered the sound signal into 128 logarithmically distributed frequency bands between 180 Hz and 7 kHz. In each band, the filtered sound signal was half-wave rectified, smoothed, and decimated to 100 Hz. The outputs of the auditory model in all frequency bands were averaged to get the envelope. The length of the time integration window D was 80, corresponding to 0.8 s. TRF(*t*) was the TRF function, which could be interpreted as the response triggered by a unit power increase of the stimulus. The TRF was independently fitted for each participant and each channel. The model was calculated using 10-fold cross-validation. Specifically, each participant’s MEG response was evenly divided into 10 segments. Nine segments were used to train the model, and the remaining segment was used to generate the prediction of neural response. The predicted neural response to the envelope was then linearly regressed out from the true neural responses, and the residual was used in the time-resolved neural encoding of time in the following.

#### Serial dependency analysis

We used multiple linear regression to predict neural activity from the current time interval (lag = 0) as well as the five preceding intervals (lags 1 to 5). For each participant, regression weights were averaged across all conditions to obtain the mean weight associated with each lag. We then tested whether the weight at each lag was significantly different from zero at the group level using one-sample t-tests with FDR correction for multiple comparisons across lags. For natural speech dataset, we exclude the syllables within the same word for the preceding intervals, because the syllable within the same word has significant correlation with the current syllable duration and therefore confounds the regression weight.

## Reference

Angeloni, C. F., Młynarski, W., Piasini, E., Williams, A. M., Wood, K. C., Garami, L., Hermundstad, A. M., & Geffen, M. N. (2023). Dynamics of cortical contrast adaptation predict perception of signals in noise. Nature Communications, 14(1), 4817. 10.1038/s41467-023-40477-6

Arnal, L. H., & Giraud, A.-L. (2012). Cortical oscillations and sensory predictions. Trends in Cognitive Sciences, 16(7), 390–398. 10.1016/j.tics.2012.05.003

Barlow, H. B. (1961). Possible principles underlying the transformation of sensory messages. Sensory Communication, 1(01), 217–233.

Barlow, H. B., & Levick, W. R. (1969). Changes in the maintained discharge with adaptation level in the cat retina. The Journal of Physiology, 202(3), 699–718. 10.1113/jphysiol.1969.sp008836

Bays, P. M., Schneegans, S., Ma, W. J., & Brady, T. F. (2024). Representation and computation in visual working memory. Nature Human Behaviour, 8(6), 1016–1034. 10.1038/s41562-024-01871-2

Benjamini, Y., & Hochberg, Y. (1995). Controlling the False Discovery Rate: A Practical and Powerful Approach to Multiple Testing. Journal of the Royal Statistical Society: Series B (Methodological), 57(1), 289–300. 10.1111/j.2517-6161.1995.tb02031.x

Bennett, R., & Elfner, E. (2019). The Syntax–Prosody Interface. Annual Review of Linguistics, 5(1), 151–171. 10.1146/annurev-linguistics-011718-012503

Blum, F., Paschen, L., Forkel, R., Fuchs, S., & Seifart, F. (2024). Consonant lengthening marks the beginning of words across a diverse sample of languages. Nature Human Behaviour, 8(11), 2127–2138. 10.1038/s41562-024-01988-4

Boersma, P. (2001). Praat, a system for doing phonetics by computer. Glot. Int., 5(9), 341–345.

Brenner, N., Bialek, W., & de Ruyter van Steveninck, R. (2000). Adaptive Rescaling Maximizes Information Transmission. Neuron, 26(3), 695–702. 10.1016/S0896-6273(00)81205-2

Brudzynski, S. M. (2026). Development and heterogeneity of rat 22 kHz vocalizations. Behavioural Brain Research, 496, 115822. 10.1016/j.bbr.2025.115822

Bu, H., Du, J., Na, X., Wu, B., & Zheng, H. (2017). AISHELL-1: An open-source Mandarin speech corpus and a speech recognition baseline. 2017 20th Conference of the Oriental Chapter of the International Coordinating Committee on Speech Databases and Speech I/O Systems and Assessment (O-COCOSDA), 1–5. 10.1109/ICSDA.2017.8384449

Buhusi, C. V., & Meck, W. H. (2005). What makes us tick? Functional and neural mechanisms of interval timing. Nature Reviews Neuroscience, 6(10), 755–765. 10.1038/nrn1764

Buzsáki, G. (2026). Time, space, memory and brain–body rhythms. Nature Reviews Neuroscience, 27(1), 61–78. 10.1038/s41583-025-00987-2

Byrd, R. H., Schnabel, R. B., & Shultz, G. A. (1988). Approximate solution of the trust region problem by minimization over two-dimensional subspaces. Mathematical Programming, 40(1), 247–263. 10.1007/BF01580735

Cao, R., Bladon, J. H., Charczynski, S. J., Hasselmo, M. E., & Howard, M. W. (2022). Internally generated time in the rodent hippocampus is logarithmically compressed. eLife, 11, e75353. 10.7554/eLife.75353

Carandini, M., & Heeger, D. J. (2012). Normalization as a canonical neural computation. Nature Reviews Neuroscience, 13(1), 51–62. 10.1038/nrn3136

Chen, G., Chai, S., Wang, G.-B., Du, J., Zhang, W.-Q., Weng, C., Su, D., Povey, D., Trmal, J., Zhang, J., Jin, M., Khudanpur, S., Watanabe, S., Zhao, S., Zou, W., Li, X., Yao, X., Wang, Y., You, Z., & Yan, Z. (2021). GigaSpeech: An Evolving, Multi-Domain ASR Corpus with 10,000 Hours of Transcribed Audio. 3670–3674. 10.21437/Interspeech.2021-1965

Cheyette, S. J., & Piantadosi, S. T. (2020). A unified account of numerosity perception. Nature Human Behaviour, 4(12), 1265–1272. 10.1038/s41562-020-00946-0

Cohen, Y. (2022). Song recordings and annotation files of 3 canaries used to evaluate training of TweetyNet models for birdsong segmentation and annotation (Version 8, p. 37829245292 bytes) [Data set]. Dryad. 10.5061/DRYAD.XGXD254F4

Corbett, J. E., Aydın, B., & Munneke, J. (2021). Adaptation to average duration. Attention, Perception, & Psychophysics, 83(3), 1190–1200. 10.3758/s13414-020-02134-8

Coupé, C., Oh, Y. M., Dediu, D., & Pellegrino, F. (2019). Different languages, similar encoding efficiency: Comparable information rates across the human communicative niche. Science Advances, 5(9), eaaw2594. 10.1126/sciadv.aaw2594

Cover, T. M., & Thomas. (1999). Elements of information theory. John Wiley & Sons.

Dean, I., Harper, N. S., & McAlpine, D. (2005). Neural population coding of sound level adapts to stimulus statistics. Nature Neuroscience, 8(12), 1684–1689. 10.1038/nn1541

Dilley, L. C., & Pitt, M. A. (2010). Altering Context Speech Rate Can Cause Words to Appear or Disappear. Psychological Science, 21(11), 1664–1670. 10.1177/0956797610384743

Ding, N., & Simon, J. Z. (2012). Emergence of neural encoding of auditory objects while listening to competing speakers. Proceedings of the National Academy of Sciences of the United States of America, 109(29), 11854–11859. 10.1073/pnas.1205381109

Doelling, K. B., Assaneo, M. F., Bevilacqua, D., Pesaran, B., & Poeppel, D. (2019). An oscillator model better predicts cortical entrainment to music. Proceedings of the National Academy of Sciences, 116(20), 10113–10121. (world). 10.1073/pnas.1816414116

Fairhall, A. L., Lewen, G. D., Bialek, W., & de Ruyter van Steveninck, R. R. (2001). Efficiency and ambiguity in an adaptive neural code. Nature, 412(6849), 787–792. 10.1038/35090500

Fitt, S. (2001). Unisyn lexicon release (Version 1.3). Edinburgh, *Scotland*: *Centre for Speech Technology Research at the University of Edinburgh*.

Frazier, L., Carlson, K., & Clifton, C. (2006). Prosodic phrasing is central to language comprehension. Trends in Cognitive Sciences, 10(6), 244–249. 10.1016/j.tics.2006.04.002

Garofolo, Lamel, Lori F., Fisher, William M., Pallett, David S., Dahlgren, Nancy L., Zue, Victor, & Fiscus, Jonathan G. (1993). TIMIT Acoustic-Phonetic Continuous Speech Corpus (p. 715776 KB) [Data set]. Linguistic Data Consortium. 10.35111/17GK-BN40

Gervain, J., & Geffen, M. N. (2019). Efficient Neural Coding in Auditory and Speech Perception. Trends in Neurosciences, 42(1), 56–65. 10.1016/j.tins.2018.09.004

Gibbon, J. (1977). Scalar expectancy theory and Weber’s law in animal timing. Psychological Review, 84(3), 279–325. 10.1037/0033-295X.84.3.279

Gibbon, J., Church, R. M., & Meck, W. H. (1984). Scalar Timing in Memory. Annals of the New York Academy of Sciences, 423(1), 52–77. 10.1111/j.1749-6632.1984.tb23417.x

Goel, A., & Buonomano, D. V. (2016). Temporal Interval Learning in Cortical Cultures Is Encoded in Intrinsic Network Dynamics. Neuron, 91(2), 320–327. 10.1016/j.neuron.2016.05.042

Grabenhorst, M., Michalareas, G., Maloney, L. T., & Poeppel, D. (2019). The anticipation of events in time. Nature Communications, 10(1), 5802. 10.1038/s41467-019-13849-0

Gramfort, A., Luessi, M., Larson, E., Engemann, D. A., Strohmeier, D., Brodbeck, C., Goj, R., Jas, M., Brooks, T., Parkkonen, L., & Hämäläinen, M. (2013). MEG and EEG data analysis with MNE-Python. Frontiers in Neuroscience, 7. 10.3389/fnins.2013.00267

Greenberg, S., Carvey, H., Hitchcock, L., & Chang, S. (2003). Temporal properties of spontaneous speech—A syllable-centric perspective. *Journal of Phonetics*, Temporal Integration in the Perception of Speech, 31(3), 465–485. 10.1016/j.wocn.2003.09.005

Hage, S. R., Gavrilov, N., Salomon, F., & Stein, A. M. (2013). Temporal vocal features suggest different call-pattern generating mechanisms in mice and bats. BMC Neuroscience, 14(1), 99. 10.1186/1471-2202-14-99

Harding, E. E., Kim, J. C., Demos, A. P., Roman, I. R., Tichko, P., Palmer, C., & Large, E. W. (2025). Musical neurodynamics. Nature Reviews Neuroscience, 26(5), 293–307. 10.1038/s41583-025-00915-4

Hayashi, M. J., & Ivry, R. B. (2020). Duration Selectivity in Right Parietal Cortex Reflects the Subjective Experience of Time. Journal of Neuroscience, 40(40), 7749–7758. 10.1523/JNEUROSCI.0078-20.2020

Hedley, R. W. (2016). Complexity, Predictability and Time Homogeneity of Syntax in the Songs of Cassin’s Vireo (Vireo cassinii). PLOS ONE, 11(4), e0150822. 10.1371/journal.pone.0150822

Heron, J., Aaen-Stockdale, C., Hotchkiss, J., Roach, N. W., McGraw, P. V., & Whitaker, D. (2011). Duration channels mediate human time perception. Proceedings of the Royal Society B: Biological Sciences, 279(1729), 690–698. 10.1098/rspb.2011.1131

Herrmann, B., Schlichting, N., & Obleser, J. (2014). Dynamic Range Adaptation to Spectral Stimulus Statistics in Human Auditory Cortex. The Journal of Neuroscience, 34(1), 327–331. 10.1523/JNEUROSCI.3974-13.2014

Huang, Q., & Doeller, C. F. (2026). Efficient coding in working memory is adapted to the structure of the environment. Cell Reports, 45(1), 116861. 10.1016/j.celrep.2025.116861

Jacewicz, E., Fox, R. A., O’Neill, C., & Salmons, J. (2009). Articulation rate across dialect, age, and gender. Language Variation and Change, 21(2), 233–256. 10.1017/S0954394509990093

Jazayeri, M., & Shadlen, M. N. (2010). Temporal context calibrates interval timing. Nature Neuroscience, 13(8), 1020–1026. 10.1038/nn.2590

Johnston, A., Arnold, D. H., & Nishida, S. (2006). Spatially Localized Distortions of Event Time. Current Biology, 16(5), 472–479. 10.1016/j.cub.2006.01.032

Katz, J., & Selkirk, E. (2011). Contrastive focus vs. discourse-new: Evidence from phonetic prominence in English. Language, 87(4), 771–816. 10.1353/lan.2011.0076

Kononowicz, T. W., & Rijn, H. van. (2014). Decoupling Interval Timing and Climbing Neural Activity: A Dissociation between CNV and N1P2 Amplitudes. Journal of Neuroscience, 34(8), 2931–2939. 10.1523/JNEUROSCI.2523-13.2014

Kopec, C. D., & Brody, C. D. (2010). Human performance on the temporal bisection task. Brain and Cognition, 74(3), 262–272. 10.1016/j.bandc.2010.08.006

Kopp-Scheinpflug, C., Sinclair, J. L., & Linden, J. F. (2018). When Sound Stops: Offset Responses in the Auditory System. Trends in Neurosciences, 41(10), 712–728. 10.1016/j.tins.2018.08.009

Kösem, A., Bosker, H. R., Takashima, A., Meyer, A., Jensen, O., & Hagoort, P. (2018). Neural Entrainment Determines the Words We Hear. Current Biology, 28(18), 2867–2875.e3. 10.1016/j.cub.2018.07.023

Kügler, F. (2008). The role of duration as a phonetic correlate of focus. 591–594. 10.21437/SpeechProsody.2008-134

Lachlan, R. F., Ratmann, O., & Nowicki, S. (2018). Cultural conformity generates extremely stable traditions in bird song. Nature Communications, 9(1), 2417. 10.1038/s41467-018-04728-1

Lakatos, P., Gross, J., & Thut, G. (2019). A New Unifying Account of the Roles of Neuronal Entrainment. Current Biology, 29(18), R890–R905. 10.1016/j.cub.2019.07.075

Landy, M. S., Trommershäuser, J., & Daw, N. D. (2012). Dynamic Estimation of Task-Relevant Variance in Movement under Risk. Journal of Neuroscience, 32(37), 12702–12711. 10.1523/JNEUROSCI.6160-11.2012

Large, E. W., & Jones, M. R. (1999). The dynamics of attending: How people track time-varying events. Psychological Review, 106(1), 119–159. 10.1037/0033-295X.106.1.119

Large, E. W., Roman, I., Kim, J. C., Cannon, J., Pazdera, J. K., Trainor, L. J., Rinzel, J., & Bose, A. (2023). Dynamic models for musical rhythm perception and coordination. Frontiers in Computational Neuroscience, 17. 10.3389/fncom.2023.1151895

Larracy, R., Phinyomark, A., Salehi, A., MacDonald, E., Kazemi, S., Bashar, S. S., Tabor, A., & Scheme, E. (2025). A dataset of high-resolution plantar pressures for gait analysis across varying footwear and walking speeds. Scientific Data, 12(1), 1415. 10.1038/s41597-025-05792-1

Laughlin, S. (1981). A simple coding procedure enhances a neuron’s information capacity. Z. Naturforsch, 36(910–912), 51.

Lewicki, M. S. (2002). Efficient coding of natural sounds. Nature Neuroscience, 5(4), 356–363. 10.1038/nn831

Lundstrom, B. N., Higgs, M. H., Spain, W. J., & Fairhall, A. L. (2008). Fractional differentiation by neocortical pyramidal neurons. Nature Neuroscience, 11(11), 1335–1342. 10.1038/nn.2212

Maslowski, M., Meyer, A. S., & Bosker, H. R. (2019). Listeners normalize speech for contextual speech rate even without an explicit recognition task. The Journal of the Acoustical Society of America, 146(1), 179–188. 10.1121/1.5116004

Mau, W., Sullivan, D. W., Kinsky, N. R., Hasselmo, M. E., Howard, M. W., & Eichenbaum, H. (2018). The Same Hippocampal CA1 Population Simultaneously Codes Temporal Information over Multiple Timescales. Current Biology, 28(10), 1499–1508.e4. 10.1016/j.cub.2018.03.051

McAuliffe, M., Socolof, M., Mihuc, S., Wagner, M., & Sonderegger, M. (2017). Montreal Forced Aligner: Trainable Text-Speech Alignment Using Kaldi. Interspeech 2017, 498–502. 10.21437/Interspeech.2017-1386

Mehr, S. A., Singh, M., Knox, D., Ketter, D. M., Pickens-Jones, D., Atwood, S., Lucas, C., Jacoby, N., Egner, A. A., Hopkins, E. J., Howard, R. M., Hartshorne, J. K., Jennings, M. V., Simson, J., Bainbridge, C. M., Pinker, S., O’Donnell, T. J., Krasnow, M. M., & Glowacki, L. (2019). Universality and diversity in human song. Science (New York, N.Y.), 366(6468), eaax0868. 10.1126/science.aax0868

Meirhaeghe, N., Sohn, H., & Jazayeri, M. (2021). A precise and adaptive neural mechanism for predictive temporal processing in the frontal cortex. Neuron, 109(18), 2995–3011.e5. 10.1016/j.neuron.2021.08.025

Mesgarani, N., David, S. V., Fritz, J. B., & Shamma, S. A. (2014). Mechanisms of noise robust representation of speech in primary auditory cortex. Proceedings of the National Academy of Sciences, 111(18), 6792–6797. 10.1073/pnas.1318017111

Młynarski, W. F., & Hermundstad, A. M. (2021). Efficient and adaptive sensory codes. Nature Neuroscience, 24(7), 998–1009. 10.1038/s41593-021-00846-0

Motanis, H., Seay, M. J., & Buonomano, D. V. (2018). Short-Term Synaptic Plasticity as a Mechanism for Sensory Timing. Trends in Neurosciences, 41(10), 701–711. 10.1016/j.tins.2018.08.001

Nicholson, D., Queen, J. E., & J. Sober, S. (2017). Bengalese Finch song repository. 10.6084/m9.figshare.4805749.v9

Nobre, A. C., & van Ede, F. (2018). Anticipated moments: Temporal structure in attention. Nature Reviews Neuroscience, 19(1), 34–48. 10.1038/nrn.2017.141

Norton, E. H., Acerbi, L., Ma, W. J., & Landy, M. S. (2019). Human online adaptation to changes in prior probability. PLOS Computational Biology, 15(7), e1006681. 10.1371/journal.pcbi.1006681

Norton, E. H., Fleming, S. M., Daw, N. D., & Landy, M. S. (2017). Suboptimal Criterion Learning in Static and Dynamic Environments. PLOS Computational Biology, 13(1), e1005304. 10.1371/journal.pcbi.1005304

Obleser, J., & Kayser, C. (2019). Neural Entrainment and Attentional Selection in the Listening Brain. Trends in Cognitive Sciences, 23(11), 913–926. 10.1016/j.tics.2019.08.004

Ofir, N., & Landau, A. N. (2022). Neural signatures of evidence accumulation in temporal decisions. Current Biology, 32(18), 4093–4100.e6. 10.1016/j.cub.2022.08.006

Oganian, Y., Kojima, K., Breska, A., Cai, C., Findlay, A., Chang, E. F., & Nagarajan, S. S. (2023). Phase Alignment of Low-Frequency Neural Activity to the Amplitude Envelope of Speech Reflects Evoked Responses to Acoustic Edges, Not Oscillatory Entrainment. Journal of Neuroscience, 43(21), 3909–3921. 10.1523/JNEUROSCI.1663-22.2023

Oldfield, R. C. (1971). The assessment and analysis of handedness: The Edinburgh inventory. Neuropsychologia, 9(1), 97–113. 10.1016/0028-3932(71)90067-4

Paschen, L., Fuchs, S., & Seifart, F. (2022). Final Lengthening and vowel length in 25 languages. Journal of Phonetics, 94, 101179. 10.1016/j.wocn.2022.101179

Passmore, S., Wood, A. L. C., Barbieri, C., Shilton, D., Daikoku, H., Atkinson, Q. D., & Savage, P. E. (2024). Global musical diversity is largely independent of linguistic and genetic histories. Nature Communications, 15(1), 3964. 10.1038/s41467-024-48113-7

Paton, J. J., & Buonomano, D. V. (2018). The Neural Basis of Timing: Distributed Mechanisms for Diverse Functions. Neuron, 98(4), 687–705. 10.1016/j.neuron.2018.03.045

Pellegrino, F., Coupe, C., & Marsico, E. (2011). A cross-language perspective on speech information rate. Language, 87(3), 539–558. 10.1353/lan.2011.0057

Pitkow, X., & Meister, M. (2012). Decorrelation and efficient coding by retinal ganglion cells. Nature Neuroscience, 15(4), 628–635. 10.1038/nn.3064

Poeppel, D., & Assaneo, M. F. (2020). Speech rhythms and their neural foundations. Nature Reviews Neuroscience, 21(6), 322–334. 10.1038/s41583-020-0304-4

Polanía, R., Woodford, M., & Ruff, C. C. (2019). Efficient coding of subjective value. Nature Neuroscience, 22(1), 134–142. 10.1038/s41593-018-0292-0

Prat-Carrabin, A., & Woodford, M. (2026). Endogenous precision of the number sense. eLife, 13, RP101277. 10.7554/eLife.101277

Qin, L., Liu, Y., Wang, J., Li, S., & Sato, Y. (2009). Neural and Behavioral Discrimination of Sound Duration by Cats. Journal of Neuroscience, 29(50), 15650–15659. 10.1523/JNEUROSCI.2442-09.2009

Rabinowitz, N. C., Willmore, B. D. B., Schnupp, J. W. H., & King, A. J. (2011). Contrast gain control in auditory cortex. Neuron, 70(6), 1178–1191. 10.1016/j.neuron.2011.04.030

Reinisch, E., & Sjerps, M. J. (2013). The uptake of spectral and temporal cues in vowel perception is rapidly influenced by context. Journal of Phonetics, 41(2), 101–116. 10.1016/j.wocn.2013.01.002

Ren, Y., Allenmark, F., Müller, H. J., & Shi, Z. (2020). Logarithmic encoding of ensemble time intervals. Scientific Reports, 10(1), 18174. 10.1038/s41598-020-75191-6

Repp, B. H., & Su, Y.-H. (2013). Sensorimotor synchronization: A review of recent research (2006–2012). Psychonomic Bulletin & Review, 20(3), 403–452. 10.3758/s13423-012-0371-2

Rimmele, J. M., Morillon, B., Poeppel, D., & Arnal, L. H. (2018). Proactive Sensing of Periodic and Aperiodic Auditory Patterns. Trends in Cognitive Sciences, 22(10), 870–882. 10.1016/j.tics.2018.08.003

Roberts, W. A. (2006). Evidence that pigeons represent both time and number on a logarithmic scale. Behavioural Processes, Proceedings of the Meeting of the Society for the Quantitative Analyses of Behavior, 72(3), 207–214. 10.1016/j.beproc.2006.03.002

Schroeder, C. E., & Lakatos, P. (2009). Low-frequency neuronal oscillations as instruments of sensory selection. Trends in Neurosciences, 32(1), 9–18. 10.1016/j.tins.2008.09.012

Schütt, H. H., Kim, D., & Ma, W. J. (2024). Reward prediction error neurons implement an efficient code for reward. Nature Neuroscience, 27(7), 1333–1339. 10.1038/s41593-024-01671-x

Schwartz, O., & Simoncelli, E. P. (2001). Natural signal statistics and sensory gain control. Nature Neuroscience, 4(8), 819–825. 10.1038/90526

Shukla, M., White, K. S., & Aslin, R. N. (2011). Prosody guides the rapid mapping of auditory word forms onto visual objects in 6-mo-old infants. Proceedings of the National Academy of Sciences, 108(15), 6038–6043. 10.1073/pnas.1017617108

Smith, E. C., & Lewicki, M. S. (2006). Efficient auditory coding. Nature, 439(7079), 978–982. 10.1038/nature04485

Soto, F., Hsiang, J.-C., Rajagopal, R., Piggott, K., Harocopos, G. J., Couch, S. M., Custer, P., Morgan, J. L., & Kerschensteiner, D. (2020). Efficient Coding by Midget and Parasol Ganglion Cells in the Human Retina. Neuron, 107(4), 656–666.e5. 10.1016/j.neuron.2020.05.030

Sun, J. Z., Wang, G. I., Goyal, V. K., & Varshney, L. R. (2012). A framework for Bayesian optimality of psychophysical laws. Journal of Mathematical Psychology, 56(6), 495–501. 10.1016/j.jmp.2012.08.002

Tsao, A., Yousefzadeh, S. A., Meck, W. H., Moser, M.-B., & Moser, E. I. (2022). The neural bases for timing of durations. Nature Reviews Neuroscience, 23(11), 646–665. 10.1038/s41583-022-00623-3

Vishne, G., Jacoby, N., Malinovitch, T., Epstein, T., Frenkel, O., & Ahissar, M. (2021). Slow update of internal representations impedes synchronization in autism. Nature Communications, 12(1), 5439. 10.1038/s41467-021-25740-y

Vuust, P., Heggli, O. A., Friston, K. J., & Kringelbach, M. L. (2022). Music in the brain. Nature Reviews. Neuroscience, 23(5), 287–305. 10.1038/s41583-022-00578-5

Wang, A. L., Mouraux, A., Liang, M., & Iannetti, G. D. (2008). The Enhancement of the N1 Wave Elicited by Sensory Stimuli Presented at Very Short Inter-Stimulus Intervals Is a General Feature across Sensory Systems. PLOS ONE, 3(12), e3929. 10.1371/journal.pone.0003929

Wang, Y., Wu, D., Ding, N., Zou, J., Lu, Y., Ma, Y., Zhang, X., Yu, W., & Wang, K. (2025). Linear phase property of speech envelope tracking response in Heschl’s gyrus and superior temporal gyrus. Cortex, 186, 1–10. 10.1016/j.cortex.2025.02.015

Weber, A. I., Krishnamurthy, K., & Fairhall, A. L. (2019). Coding Principles in Adaptation. Annual Review of Vision Science, 5(Volume 5, 2019), 427–449. 10.1146/annurev-vision-091718-014818

Wiener, M., & Thompson, J. C. (2015). Repetition enhancement and memory effects for duration. NeuroImage, 113, 268–278. 10.1016/j.neuroimage.2015.03.054

Willmore, B. D. B., & King, A. J. (2023). Adaptation in auditory processing. Physiological Reviews, 103(2), 1025–1058. 10.1152/physrev.00011.2022

Zada, Z., Nastase, S. A., Aubrey, B., Jalon, I., Michelmann, S., Wang, H., Hasenfratz, L., Doyle, W., Friedman, D., Dugan, P., Melloni, L., Devore, S., Flinker, A., Devinsky, O., Goldstein, A., & Hasson, U. (2025). The “Podcast” ECoG dataset for modeling neural activity during natural language comprehension. Scientific Data, 12(1), 1135. 10.1038/s41597-025-05462-2

Zalta, A., Large, E. W., Schön, D., & Morillon, B. (2024). Neural dynamics of predictive timing and motor engagement in music listening. Science Advances, 10(10), eadi2525. 10.1126/sciadv.adi2525

Zhang, B., Lv, H., Guo, P., Shao, Q., Yang, C., Xie, L., Xu, X., Bu, H., Chen, X., Zeng, C., Wu, D., & Peng, Z. (2022). WENETSPEECH: A 10000+ Hours Multi-Domain Mandarin Corpus for Speech Recognition. ICASSP 2022 - 2022 IEEE International Conference on Acoustics, Speech and Signal Processing (ICASSP), 6182–6186. 10.1109/ICASSP43922.2022.9746682

Zhang, Y., Leonard, M. K., Bhaya-Grossman, I., Gwilliams, L., & Chang, E. F. (2026). Human cortical dynamics of auditory word form encoding. Neuron, 114(1), 167–180.e6. 10.1016/j.neuron.2025.10.011

Zhang, Y., Pan, C., Guo, W., Li, R., Zhu, Z., Wang, J., Xu, W., Lu, J., Hong, Z., Wang, C., Zhang, L., He, J., Jiang, Z., Chen, Y., Yang, C., Zhou, J., Cheng, X., & Zhao, Z. (2024). GTSinger: A Global Multi-Technique Singing Corpus with Realistic Music Scores for All Singing Tasks. Advances in Neural Information Processing Systems, 37, 1117–1140. 10.52202/079017-0034

Zhang, Y., Zou, J., & Ding, N. (2023). Acoustic correlates of the syllabic rhythm of speech: Modulation spectrum or local features of the temporal envelope. Neuroscience & Biobehavioral Reviews, 147, 105111. 10.1016/j.neubiorev.2023.105111

Zoefel, B., Archer-Boyd, A., & Davis, M. H. (2018). Phase Entrainment of Brain Oscillations Causally Modulates Neural Responses to Intelligible Speech. Current Biology, 28(3), 401–408.e5. 10.1016/j.cub.2017.11.071

Zou, J., Xu, C., Luo, C., Jin, P., Gao, J., Li, J., Gao, J., Ding, N., & Luo, B. (2021). θ-Band Cortical Tracking of the Speech Envelope Shows the Linear Phase Property. eNeuro, 8(4). 10.1523/ENEURO.0058-21.2021

